# Patterns of piRNA regulation in *Drosophila* revealed through transposable element clade inference

**DOI:** 10.1101/2021.04.29.442051

**Authors:** Iskander Said, Michael P. McGurk, Andrew G. Clark, Daniel A. Barbash

## Abstract

Transposable elements (TEs) are self-replicating “genetic parasites” ubiquitous to eukaryotic genomes. In addition to conflict between TEs and their host genomes, TEs of the same family are in competition with each other. They compete for the same genomic niches while experiencing the same regime of copy-number selection. This suggests that competition among TEs may favor the emergence of new variants that can outcompete their ancestral forms. To investigate the sequence evolution of TEs, we developed a method to infer clades: collections of TEs that share SNP variants and represent distinct TE family lineages. We applied this method to a panel of 85 *Drosophila melanogaster* genomes and found that the genetic variation of several TE families shows significant population structure that arises from the population-specific expansions of single clades. We used population genetic theory to classify these clades into younger versus older clades and found that younger clades are associated with a greater abundance of sense and antisense piRNAs per copy than older ones. Further, we find that the abundance of younger, but not older clades, is positively correlated with antisense piRNA production, suggesting a general pattern where hosts preferentially produce antisense piRNAs from recently active TE variants. Together these findings suggest a pattern whereby new TE variants arise by mutation and then increase in copy number, followed by the host producing antisense piRNAs that may be used to silence these emerging variants.

## Introduction

Transposable elements (TEs) are mobile, selfish genetic elements commonly thought of as “genetic parasites’’. At the start of an invasion TEs begin as a single copy within a host genome, but can transpose and expand rapidly in copy number throughout the population in each successive generation by using the host’s replication machinery (Doolittle and Sapienza 1980; Orgel and Crick 1980). In *Drosophila,* explosive growth in copy number during a TE invasion is thought to be quickly followed by the host acquiring resistance to TE transpositions, commonly through host production of *piwi*-interacting small RNAs, piRNAs which interfere with TE transcripts (Le Rouzic and Capy 2005; Aravin et al. 2007; Brennecke et al. 2007; Kofler et al. 2018). In the germline, piRNA-mediated silencing is established through the activation of two synergistic pathways. In the “primary” piRNA pathway, transcription of long precursor sense and antisense RNAs from TE-rich loci called piRNA clusters are processed into 21-30 base-pair-long antisense piRNAs that complex with Piwi clade proteins, bind to nascent sense TE transcripts by recognizing sequence complementarity, and then recruit additional proteins to transcriptionally silence homologous TEs. In the “secondary” piRNA pathway, Piwi*-*bound piRNAs degrade TE transcripts and form sense piRNAs that bind to antisense piRNA precursors and create a positive feedback loop, known as the Ping-Pong cycle, that establishes constitutive silencing (Aravin et al. 2007; Brennecke et al. 2007; Le Thomas et al. 2014; Czech et al. 2018).

Ultimately the TE copy number may reach a steady state, with the rate of transposition dampened by piRNA silencing as well as selection against the deleterious consequences to reproductive fitness of the host organism (Charlesworth and Charlesworth 1983; Lee and Langley 2010; Kelleher et al. 2020). However, as TEs expand in copy number, they also acquire polymorphisms in their sequences, which may lead to the formation of new lineages or subfamilies (Moody 1988; Kimmel and Mathaes 2010; Kijima and Innan 2013; Iwasaki et al. 2020). Multiple lineages of a TE will compete with each other, as long as their polymorphisms are not deactivating (Abrusán and Krambeck 2006; Le Rouzic and Capy 2006; Iwasaki et al. 2020). Much in the same way individuals within an ecological system are constrained by a carrying capacity, variants of the same TE may be constrained by the copy-number carrying capacity of the host (Brookfield 2005; Le Rouzic and Capy 2006). This dynamic produces an arena of genomic competition of TE variants where selection may drive the propagation of more fit TE lineages, while less fit lineages are purged (Le Rouzic and Capy 2006).

The study of selection on TE population variation has often focused on the fitness of the host organism rather than on the TEs themselves. Much of it centered on the variation of TE insertions within and between populations as well as fitness and phenotypic effects associated with particular insertion loci (Cridland et al. 2013; Blumenstiel et al. 2014; Kofler et al. 2015). The study of selection on sequence variation of TEs, on the other hand, is much more limited. TEs are typically categorized into classes and subclasses based first on their mechanism of transposition, and then on presence of shared motifs, relative sequence identity, and phylogenetic characteristics (Wicker et al. 2007; Arkhipova 2017; Makałowski et al. 2019). There is extensive systemization of TE families, describing their consensus sequences, open reading frames, and insertion site preferences (Bao et al. 2015).

Due to the challenges of quantifying variation within repetitive sequences, however, the empirical study of TE sequence polymorphism is largely limited to analyses of reference genome assemblies. For example, the sequence variation of TE families in the *Drosophila melanogaster* reference genome has been comprehensively described (Kaminker et al. 2002; Lerat et al. 2003; Bergman and Bensasson 2007). In another example, phylogenetic and evolutionary analyses on retrotransposons within the *Oryza sativa* genome revealed strong purifying selection on protein-coding regions, with occasional bursts of positive selection (Baucom et al. 2009). To our knowledge, studies examining TE sequence variation from population samples are rare, likely because reference genomes are the primary source of full-length TE sequence data.

Ideally, to apply population genetic and molecular evolutionary principles to the genetic variation of TEs, we would study the complete sequences of individual TE insertions across many genomes. This is especially necessary if the aim is to assess competition between TE subfamily lineages, where reconstructing the underlying phylogenies would yield insight into the dynamics of how lineages diversify and potentially compete with each other. However, most current population genomic data comes from short-read sequencing, which does not permit an unambiguous assembly containing all the TEs and their internal SNPs. The problem is related to haplotype phasing, which can be done with short reads (Clark 1990; Excoffier and Slatkin 1995; Browning and Browning 2007; Delaneau et al. 2008), except here the TE insertions are at nonhomologous positions. Furthermore, the high multiplicity of TEs greatly complicates the task of determining which internal polymorphisms co-occur in the same insertion, such that with short reads, unambiguous TE haplotypes cannot be recovered as complete sequences of linked SNPs. Although new long-read technologies, like PacBio and Oxford Nanopore, have emerged that greatly reduce phasing problems and allow for more complete analysis of TEs (Long et al. 2018; Miller et al. 2018; Chakraborty et al. 2019; Ellison and Cao 2020; Wierzbicki et al. 2020; Gebert et al. 2021 Jul 30), their higher cost and relatively high error rates have limited their application to large-scale population genomics studies.

To gain insight into the sequence evolution of actively invading TEs, we sought to resolve some of these challenges by leveraging a straightforward intuition: If a set of SNPs co-occur in the same TE lineage, their copy-number variation should be correlated across genomes, covarying as the copy number of that lineage varies across genomes. To this end, we took advantage of the large sample sizes in a population-genomic dataset to quantify positive correlations in the copy number of SNPs across multiple individuals, and from these we identify groups of SNPs that we infer to co-occur within TE lineages. We refer to these groups of SNPs as clades, which are inferred to distinguish lineages of TE subfamilies while sidestepping the task of reconstructing the full phylogenies from short-read data. We use the term “TE lineages” to describe the true but unknown genealogical relationship of TE sequences in a family, while “clades” are statistical inferences of this genealogy informed by the positive correlation of SNP copy number. Applying our method to a set of 85 *D. melanogaster* genomes from the Global Diversity Lines (GDL) (Grenier et al. 2015), we inferred clades in 41 recently active TE families. We then used public PacBio datasets and simulations to validate our inferred clades (Long et al. 2018; Chakraborty et al. 2019). We analyzed the population variation of TE variants and found significant population structure driven by population-specific TE clades, several which are likely active. We additionally analyzed several piRNA libraries from ovaries focused on SNPs that distinguish clades, and found piRNAs are especially enriched for younger TE clades.

## New approaches

### Hierarchical clustering of SNPs uncovers TE clades in NGS data

We leverage large population-genomic datasets to detect TE clades by inferring the co-occurrences of SNPs within a TE lineage. We expect that if two alleles exist within the same lineage they will correlate in copy number, varying together as the TEs of that lineage vary in copy number (Figure 1a). We apply this principle to all pairwise combinations of SNPs within an element to compute a correlation matrix and then use hierarchical clustering to cluster groups of SNPs that are strongly correlated. The result is clusters of SNPs that co-vary in their copy number across samples; because these are inferred to occur within the same TE lineage we refer to these clusters as “clades”. Hierarchical clustering is a particularly appropriate choice for this problem as SNPs within TE lineages are truly related to each other in an underlying tree-like structure that is analogous to a hierarchical clustering dendrogram. The correlations between alleles are unlikely to be a result of co-transposition of multiple TEs because the linkage between TEs is very low and the sampling variation in these data is quite high.

**Figure 1.**
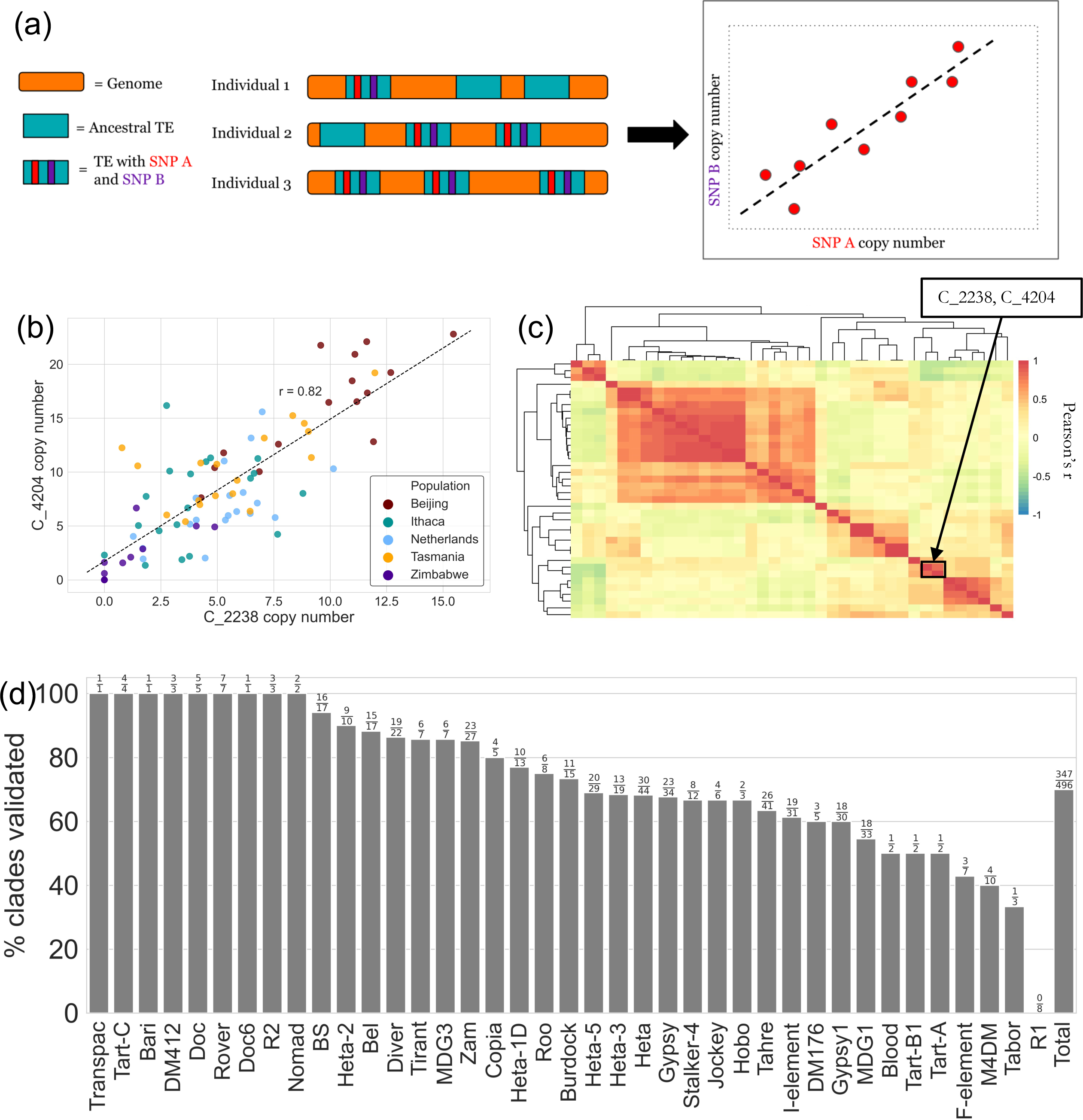
Outline and examples of the TE clade inference method. (a) Cartoon depicting the method. The genomes of three individuals (orange rectangles) contain copies of an ancestral TE (blue rectangles) and a derived TE with SNPs A and B (blue rectangles with a red and purple stripe, respectively). As the copy number of the derived TE varies in copy number across individuals 1-3, so does the copy number of SNPs A and B. This relationship in copy number is depicted as a cartoon scatterplot, where each red dot represents the copy number of SNPs A and B in one of nine individuals sampled in the population. The copy number of the SNPs is positively correlated because the SNPs are physically linked. (b) Scatterplot depicting the correlation in copy number across GDL individuals for two SNPs in the *Jockey* element. Each dot represents the copy number of the SNPs C at position 2238 (C_2238) and C at position 2402 (C_4204) for each individual, colored by their population of origin. The degree of correlation of these two SNPs is high (Pearson’s *r = 0.82*), suggesting that they are physically linked and represent a clade. Black dashed line is a linear fit of the data drawn for emphasis. (c) Heatmap showing correlation of the copy number of all SNPs from the *Jockey* element. Cells in the heatmap are seriated via hierarchical clustering to create clusters of tightly correlated SNPs, which are inferred to be *Jockey* clades segregating in the population. The cells are shaded by the pairwise Pearson’s correlation between SNP copy number. The SNPs from (b) are outlined in a block box. (d) The percent of clades inferred from GDL data that were then detected in a set of PacBio genomes (includes only clades where at least two SNPs were detected at any frequency in the PacBio data). The results are separated by TE family, with total clades shown on the far right. Fraction of clades validated over the total number of clades found are placed above each bar.

We employed this clade inference method using short-read libraries from the Global Diversity Lines (GDL), 85 *D. melanogaster* lines from populations in Beijing, Ithaca, the Netherlands, Tasmania, and Zimbabwe (Grenier et al. 2015). We aligned the short-read data to the TE consensus sequences of 41 recently active TEs and the *D. melanogaster* reference genome using ConTExt (McGurk and Barbash 2018). Then we calculated allele frequencies of SNPs from the read pileups of the alignments and calculated copy number from the read depth. In brief, copy number was estimated by dividing the observed read depth at each position on the TE consensus sequence by the expected read depth of single copy sequences inferred from the read depth of the reference genome, with corrections for GC bias (McGurk et al. 2020 Dec 22). For each individual we took the allele frequencies at each position and multiplied them by the estimated copy number at each position to generate the copy number of alleles at each position for that individual. We compute pairwise correlations between the copy number of alleles across individuals (Figure 1b), and then employ hierarchical clustering to cluster positively correlated SNPs, thus inferring clades (Figure 1c). For each of the 41 TEs analyzed, we report the SNP clusters, as well as the copy number of each inferred clade, calculated by averaging the copy number of the individual alleles (*e.g.* C_2238, C_4204 in Figure 1) (Supplemental File 1, https://github.com/is-the-biologist/TE_CladeInference).

One important consideration of this method is that we are not identifying the full set of TE insertions within an inferred clade nor the complete sequences of the insertions at any particular locus that belong to an inferred clade. Rather, we are identifying sets of SNPs that distinguish lineages from each other (lineage-informative SNPs). Lineage-informative SNPs are clustered together by our method and output as inferred clades, which are statistical inferences of the set of true lineages that exist for a TE family. However, because all TE lineages of a family are related by an underlying phylogeny, there likely does not exist a single correlation cut- off that optimally groups TEs into distinct clades. Rather, any chosen threshold induces some degree of coarse-graining in how it collapses this phylogeny, splitting and merging lineages of TE variants into clades. Generally, a higher stringency in the clustering cut-off will produce many small clusters of tightly correlated SNPs that split lineages, while low stringency cut-offs will produce a few large clusters that merge lineages together. By analyzing the phylogeny of full length TE insertions for a given family, we found that insertions that belong to clades inferred under stringent clustering parameters tend to be more closely related phylogenetically than those inferred under lenient clustering parameters (Supplemental Figure 1). Therefore, the degree of correlation between the copy number of alleles is related to the phylogenetic distance between the TE insertions that bear those alleles. However, this relationship is not always perfect. For example, a pair of alleles that are present in all TE insertions in a population may be strongly correlated, but the phylogenetic distance between TE insertions bearing those alleles may be relatively large, as for “Cluster_7’’ of the *Jockey* element (Supplemental Figure 1d, 1e).

Due to the inherent coarse-graining of hierarchical clustering, whether or not SNPs are merged into one clade depends on the clustering cut-off and how often the lineage-informative SNPs co-occur. This can result in an insertion belonging to more than one clade, depending on the correlation cut-off chosen. For example, two closely related lineages may share an ancestral set of SNPs, but have recently diverged such that one lineage acquired a small number of polymorphisms in addition to the ancestral SNPs. Depending on the stringency of the clustering cut-off these two lineages may be called as a single clade containing all of the SNPs or as two clades: one only with the ancestral subset of SNPs and one only with the derived SNPs. In the latter case of two clades, TE insertions from the derived lineage have both the ancestral and derived SNPs and would be classified as belonging to both clades. Therefore, clades should not necessarily be interpreted as distinct waves of invasion nor as being collections of unique insertions, but rather as clusters of SNPs that co-occur within insertions that are statistical representations of evolving lineages.

We attempt to address these caveats and trade-offs in our analysis by using simulations and PacBio data to validate our inferences. When inferring clades in the GDL short-read data, we chose stringent clustering parameters to make the clade calls conservative (splitting distantly related lineages). These sets of parameters were chosen to essentialize TE clades to a minimum number of core SNPs that co-occur with a high positive correlation, while increasing the number of distinct, resolved clades. High stringency cut-offs also minimize the number of false positives that can result from performing many thousand pairwise correlations of allele copy number (Supplemental Figure 1a, 1b). Parameters could be tuned to be less stringent to define clades harboring greater internal SNP variation, but the sets of parameters we chose performed well in our validation as 70% of detectable clades inferred from the GDL data were detected in long-read PacBio assemblies (Figure 1d). Undetected clades were a mix of closely related lineages that had been merged into a single cluster and errors in clustering (see *Methods*). Our method performed well on simulated datasets under a wide range of clustering parameters and under various degrees of simulated sequencing error, but was most negatively affected by excessive fragmentation of elements (Supplemental Figure 2). We additionally assessed the robustness of our method to smaller sample sizes by down-sampling the GDL data and found that when sample sizes were below 50 libraries the ability to recover accurate cluster labels dropped, especially in LINE-like retrotransposons (Supplemental Figure 3a).

## Results

### Diversity and variation of TE clades in the GDL

To determine whether high sequence diversity of a TE family is due to the evolution of many distinct lineages or a few highly diverged lineages, we assessed the sequence diversity of TE families and the number of clades segregating in the GDL. We calculated the average nucleotide diversity across the GDL of active TE families and found a positive relationship between the number of clades and nucleotide diversity (Figure 2a; Pearson’s *r = 0.74; p-value < 0.05*). Both the telomeric TEs and LTR retrotransposons have several families with a high number of clades and high sequence diversity. This observation conflicts with previous reports of LTR retrotransposons being typically less diverse than other classes of transposons because they are younger and had recently invaded *D. melanogaster* (Bergman and Bensasson 2007). Our higher estimates of sequence diversity may be a result of our sampling population-level data as opposed to previous studies that were limited to a single reference genome. It should also be noted that the LTR families *Zam, Gypsy* and *Gypsy1* are the most diverse, have the highest number of clades and are thought to be older than other LTR families in our list (Kofler et al. 2012; Kelleher and Barbash 2013; Kofler et al. 2015). Due to their age, their high diversity likely reflects inactive and degenerate lineages. However, nucleotide diversity is also a product of the mutational processes of a TE, and differences in sequence diversity between classes may also be driven by differences in the rate of mutation.

**Figure 2.**
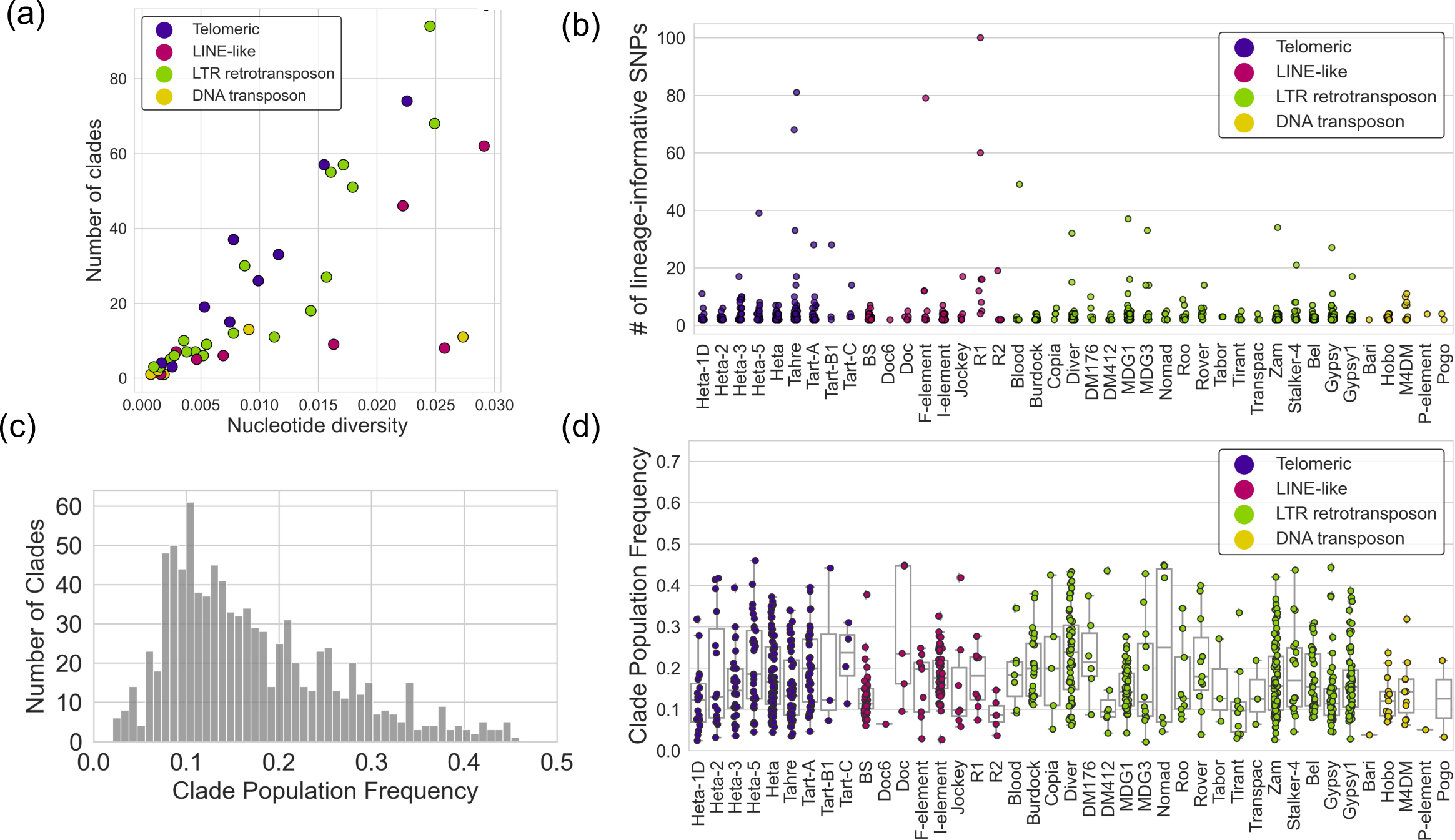
Summary statistics of TE clades inferred from GDL short-reads. (a) Average nucleotide diversity for each TE family vs. the number of clades inferred for that family, colored by TE class. (b) The number of phase-informative SNPs that compose each clade inferred for every TE family. Each point represents a clade and is colored by TE class. (c) Histogram of the clade population frequency of all clades from 41 recently active TE families. (d) Boxplots of the clade population frequency of all clades separated by TE family. Each point represents a clade and is colored by TE class.

Besides the LTR retrotransposons, there are also several LINE-like retrotransposon and DNA transposon families with high sequence diversity, but they tend to have fewer clades than non-LINE retrotransposons with similar sequence diversity. For example, the *R1* family has a high sequence diversity (*π* ≈ 0.026) and only eight clades, while the LTR retrotransposon *Zam* has comparable sequence diversity and 94 clades. This pattern may be driven by merging SNPs into large clades rather than splitting them into many small ones, so we characterized each clade by the number of lineage-informative SNPs it contains (Figure 2b). In general, the distribution of lineage-informative SNPs is small and tightly distributed, with a median number of two and an interquartile range (IQR) of one. The small cluster size is indicative of a preference for splitting multiple related clades rather than merging them into larger clusters. Clusters of SNPs we discover may therefore not be mutually exclusive and may occur together within a subset of insertions, but with a degree of positive correlation insufficient to pass the clustering threshold. This preference for splitting would upwardly bias the number of clades that we estimate, as large clusters of SNPs with distant genealogical relationships might be broken up into many small clusters of SNPs that are closely related. The exact number of clades segregating in each TE family is affected by parameterization of the clustering as well as the degree of fragmentation of the TE in the genomes themselves. Therefore, some caution must be taken when considering the absolute numbers of clades segregating in a TE family and not relative proportions. However, the inference of clades using simulated data shows that the number of clades and quality of clustering is surprisingly robust to clustering parameters (Supplemental Figure 2).

Although most TE families have clades with a small number of SNPs, *R1* clades are notable outliers, with a median of 14 SNPs and IQR 19.75 and two clades with 60 and 100 SNPs each. This might be explained by the presence of two independently evolving populations of *R1* elements in *D. melanogaster*, the hundreds of *R1* insertions in the highly repetitive ribosomal DNA array, and a separate lineage of divergent elements that comprise a megabase-sized satellite array (Wellauer and Dawid 1977; Roiha et al. 1981; Xiong and Eickbush 1988; Luan et al. 1993; McGurk and Barbash 2018). Divergence between these lineages likely explains the high sequence diversity. The similarity of sequences within each lineage and dissimilarity between lineages may favor their merging into a handful of large clades during clustering.

We next addressed whether a single clade dominates a transposable element family in terms of copy number or if instead the clades occur at roughly equal frequency. We determined the proportion of all copies of a TE family in the GDL population belonging to a clade by calculating the “clade population frequency”, dividing the total clade copy number by the average copy number across the lineage-informative positions (*e.g.* position 2238, position 4204 in Figure 1) summed across the GDL (Supplemental File 1, https://github.com/is-the-biologist/TE_CladeInference). It should be noted that clade population frequency in this context is not the frequency of any insertion of a TE in the population, but the number of copies belonging to a clade out of all copies in a population. We find that most clades occur in the population at a frequency between ∼10-30% (mean 17%; Figure 2c). There is a notable lack of low frequency clades, likely due to the filtering of low copy-number and low population-frequency alleles before calling clades. We found 21 clades with high population frequency ( > 40%) occurring in 13 different TE families, including telomeric TEs and several LTR and LINE-like retrotransposons. The *Diver* and *Nomad* LTR retrotransposons had the greatest number of high frequency clades (3-4 per family), but with dramatically different clade population frequency distributions. The *Diver* frequency distribution had many clades spread across the entire range from low to high, while the *Nomad* distribution was clearly split between a handful of low and high frequency clades (Figure 2d). *Jockey, Doc*, and *DM412* are examples of TE families with *Nomad*-like frequency distributions, while *Gypsy1*, *Zam*, and *I-element* are examples of TE families with more uniform clade frequency distribution similar to *Diver* (Figure 2d). The *Nomad- like* frequency distributions may reflect a relatively fast copy-number expansion of a handful of clades that outcompeted other lineages, while *Diver-like* distributions may reflect gradual diversification and slow increase in copy number of many clades, possibly driven by stochastic processes.

One important consideration is that due to the way population frequency was calculated, clades with SNPs in commonly deleted portions of a TE may be at a high frequency despite being at a relatively low copy number. This is particularly important for LINE-like elements and DNA transposons that are frequently truncated and internally deleted. Therefore, the clade population frequency does not necessarily reflect the proportion of TE insertions in the clade, but instead the number of TE insertions that have those nucleotide sites in the given clade. High population-frequency clades in *Gypsy*, *Zam*, *HeT-A2*, and *Tart-B1* had very low copy numbers (∼1-2 copies on average; Supplemental File 1, https://github.com/is-the-biologist/TE_CladeInference), likely due to having SNPs in commonly deleted portions of their respective TE sequences. However, we found that many high population-frequency clades were at high copy number (10-40 copies on average). The high frequency clades of *Jockey, DM412*, and *Doc* were particularly striking as they are at high copy number and dominate other clades of their respective families. These clades may be at high frequency due to age, having a competitive edge over other variants, or by pure chance.

### The majority of clades are young and recently active

The frequency of individual TE insertions in a population, or “insertion-site frequency”, is an informative parameter in estimating the age and potential activity of TEs. While the limitations of short-read data prevent us from mapping SNPs to specific insertions and estimating insertion-site frequency, the insertion-site frequency spectrum of a TE influences the variance of its copy-number distribution across individuals. Our approach infers TE clades, which are collections of TE insertions that share a subset of SNPs, but the same idea applies -- the insertion-site frequency spectrum, and therefore copy-number distribution, of clades is also expected to be related to their age. We therefore applied population genetic theory to predict the age of clades from their copy-number distributions. The copy-number distribution of a transposable element lineage in a population is related to the insertion-site frequency spectrum:

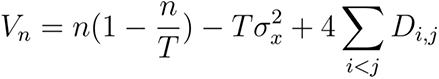

Where, *V_n_* and *n* are the variance and the mean of the copy-number distribution, respectively. *T* is the number of occupiable sites of a TE, *σ*_x_^2^ is the variance of the insertion-site frequency, and 4∑*D_i,j_* represents the sum of the coefficients of linkage disequilibrium between insertions (Charlesworth and Charlesworth 1983; Langley et al. 1983).

In a simple scenario, a recently active, young lineage will have insertions that are mostly at a low population frequency with little variance (*σ*_x_^2^ ≅ 0). When the number of occupiable sites is large (*T* >> *n*) and the effect of linkage disequilibrium between insertions is small (4∑*D_i,j_* ≅0), then the mean and variance of the copy-number distribution will be Poisson distributed:

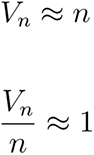

An older lineage, on the other hand, will have insertions at variable frequencies (*σ*_x_^2^ > 0) -- due to drift increasing the frequency of older insertions. This results in the variance of the copy-number distribution being less than the mean *i.e.* “underdispersed” relative to the Poisson expectation:

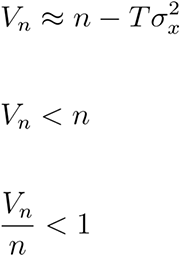

Therefore, an underdispersed copy-number distribution is indicative of a lineage with a broad insertion-site frequency spectrum, which would indicate older age and inactivity (Charlesworth and Charlesworth 1983; Langley et al. 1983; McGurk et al. 2020 Dec 22).

This is further complicated by linkage disequilibrium between insertions (*e.g.* population structure). For a recently active lineage (*σ*_x_^2^ ≅ 0) with population structure, the copy-number distribution will no longer be Poisson. Instead, the variance of the copy number will be greater than the mean *i.e.* “overdispersed”:

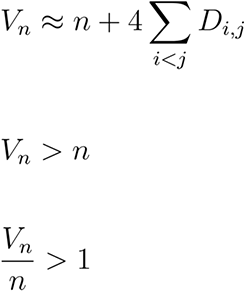

It is possible that an older TE lineage (*σ*_x_^2^ > 0) could be experiencing population structure as well, in which case whether or not the copy number distribution remains underdispersed depends on how strong the linkage is between insertions. More generally, when *Tσ*_x_^2^ > 4∑*D_i,j_* the TE will be underdispersed, when *Tσ*_x_^2^ < 4∑*D_i,j_* the TE will be overdispersed and when *Tσ*_x_^2^ ≅ 4∑*D_i,j_* the TE will fit a Poisson (Charlesworth and Charlesworth 1983; Langley et al. 1983; McGurk et al. 2020 Dec 22). Although the theoretical expectation is that young lineages will fit a Poisson and old lineages will be underdispersed, whether these expectations are borne out in the data is dependent on the population structure and demographic history of the organism. Further complicating our expectations are the complex life histories of TEs, such as recurrent invasions or extended continuous activity, wherein ancient TE insertions would be at high frequency but recent insertions would be at low frequency, thus creating an “underdispersed” copy number distribution despite having recent transpositions.

However, in practice the copy-number distributions of known active and inactive TE families recapitulate these expectations well (McGurk et al. 2020 Dec 22). Therefore, we analyzed the copy-number distribution of clades (as a proxy for the true lineages) by using a two-tailed dispersion test with multiple testing correction to ask whether the distributions are overdispersed, underdispersed, or fit a Poisson (Figure 3a) (Yang et al. 2009). We consider clades that have copy-number distributions that fit a Poisson or are overdispersed to be younger, recently active lineages and clades that are underdispersed to be older, likely inactive lineages. By using this test we are not directly assaying transpositional activity of these clades, but are merely inferring their age based on the theoretical expectations of their copy-number distribution. However, due to the caveats mentioned above we sought to validate our inferences of young and old clades by finding insertions belonging to these clades in the PacBio data and determining their sequence diversity, length, and genomic position. We found that insertions belonging to young clades tend to have much lower sequence diversity (***π*** = ∼0.05), are more often full length and are found less often in heterochromatin than insertions belonging to old clades (***π*** = ∼0.16) (*Mann-Whitney U: p-value* < 0.05; Supplemental Figure 4). These results confirm that clades with underdispersed copy-number distributions are composed of older, more degenerate insertions that have accumulated in the heterochromatin and match our theoretical expectation.

**Figure 3.**
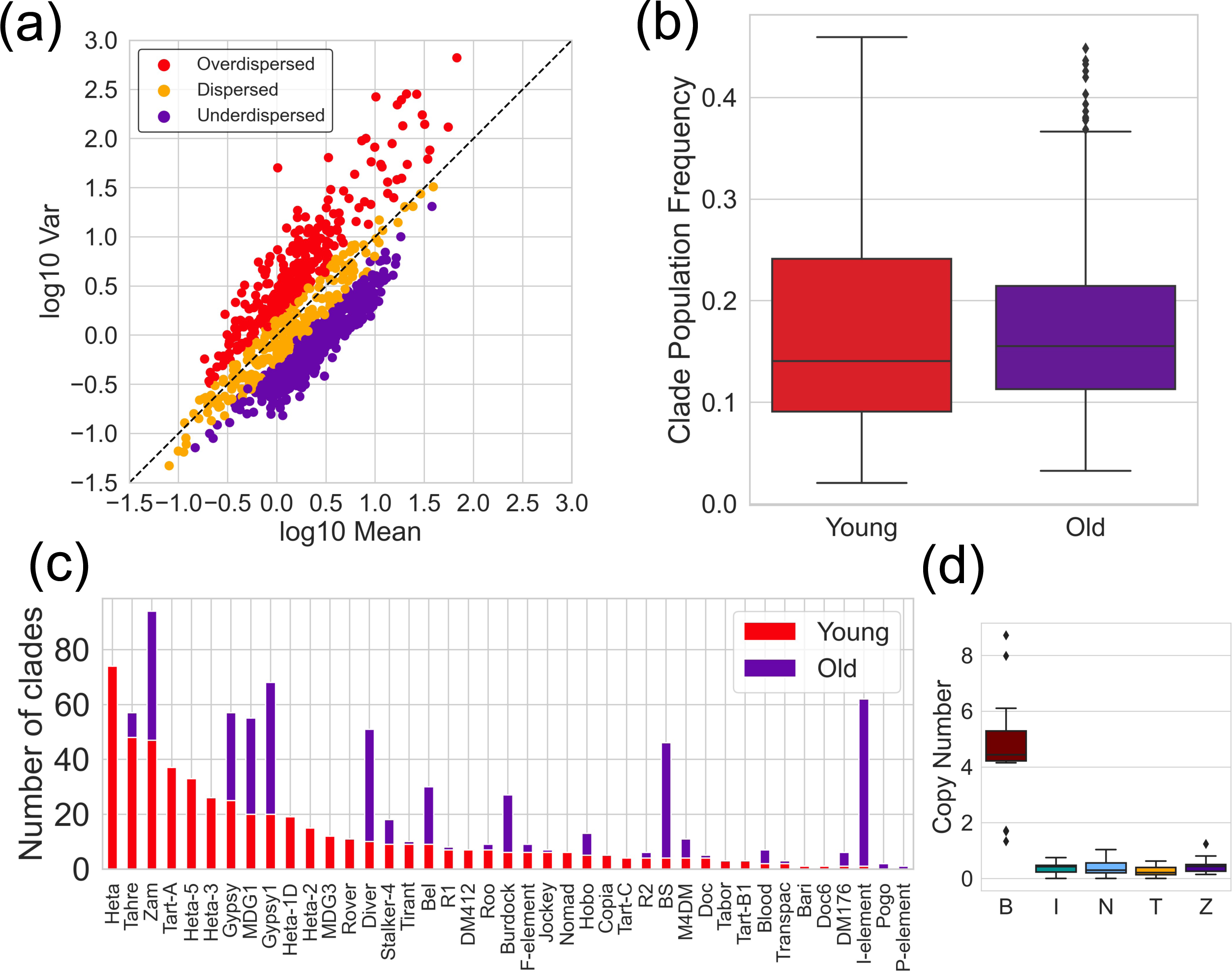
Age of clades are inferred by their copy-number distribution. (a) Mean-variance relationship of the clade copy-number distributions for clades from all families. The copy-number distributions for each clade were tested for goodness of fit to a Poisson distribution, and then colored based on acceptance or rejection of this test: “overdispersed” (rejected, red), “dispersed” (fail to reject, yellow), or “underdispersed” (rejected, purple). (b) Clade population frequency of young (red: “dispersed’’ and “overdispersed”) or old (purple: “underdispersed”) clades across all TE families. (c) Number of discovered clades per TE family that are young (red: “dispersed’’ and “overdispersed”) or old (purple: “underdispersed”). (d) Boxplot of the copy-number distribution of the sole putatively active *I-element* clade from (c) for each GDL population. There is a significant elevation in the copy number of this clade in Beijing.

Of the clades that were at a high clade population frequency (> 40%), the clades of telomeric TEs and other families such as *Jockey, Copia, Nomad* and *DM412* had copy-number distributions consistent with recent activity (dispersed or overdispersed), while the high frequency clades of *Zam* and *Stalker-4* were classified as older lineages (underdispersed). High frequency clades in *Doc* and *Diver*, on the other hand, were a mixture of young and old clades. We found that on average old clades were at a slightly higher frequency in the population than young clades (*Mann-Whitney U: p-value < 0.05;* Figure 3b). Overall, 56% of the clades were classified as young and therefore active. The excess of old clades segregating in the GDL is driven by eight families (*Gypsy*, *Gypsy1*, *I-element*, *BS*, *Zam, Bel*, *Diver* and *Burdock*), which accounts for 86% of all the old clades (Figure 3c). The abundance of old clades in these TE families matches their known insertion-site frequency spectra, which is skewed towards older insertions at high frequencies (Kofler et al. 2012; Kofler et al. 2015). Curiously, only a single old clade was inferred for the *P-element* family (Figure 3c). *P-element* is not an old TE family, but nonetheless this clade is underdispersed and at very low copy number (∼0.4 copies on average). This suggests that this clade of *P-element* is dead and possibly internally deleted. This does not mean that all *P-element* lineages are inactive, but it is the only *P-element* variant segregating in the GDL that met our criterion of detectability. Most active *P-elements* will be most similar to the consensus TE sequence, because it has very low genetic diversity (***π*** = ∼0.00076). This is also likely the case for *Pogo* where a similar pattern is seen.

*I-element*, by contrast, is a particularly striking example where all but one clade is old (Figure 3c). This is consistent with the known evolutionary history of the *I-element*, as it appears to have invaded *D. melanogaster* populations multiple times, leaving both inactive relics of ancient invasions and younger active copies (Picard et al. 1978; Kidwell 1983; Busseau et al. 1994). Many *D. melanogaster* strains are susceptible to *I-element* invasion, despite having euchromatic insertions, and crosses with strains carrying active *I-elements* result in hybrid dysgenesis (Olovnikov et al. 2013; Ryazansky et al. 2017). Therefore, many of the older clades segregating in *I-element* may be remnants of this ancient invasion.

Curiously, the only *I-element* clade that was predicted to be young shows strong population structure, being at a higher copy number in Beijing than in the other populations (Figure 3d). Strong population structure is expected to inflate the variance in copy number calculated across all populations, so we re-analyzed the *I-element* in the Beijing population. We found the copy-number distribution of the Beijing strains fit a Poisson well, implying that this *I-element* clade is likely young and active (*p-value* = 0.7). This clade is at less than 1% frequency in the other four populations, while at ∼11% frequency in the Beijing population. It is quite likely then that this young *I-element* clade invaded the Beijing population after the *D. melanogaster* population migrated to East Asia. Generally, 54% of the likely active clades were overdispersed, which suggests there may be population structure to their geographic distributions as well and potentially indicates ongoing population-specific invasions. Alternatively, overdispersion may be driven by ongoing copy-number evolution of the TE families in the strains maintained in the laboratory, thus creating an extreme form of population structure.

### Population structure of TE variation

The genetic variation of TEs within and between populations is an underexplored facet of TE evolution. Early in the *P*-element and *hobo* invasions, variant lineages emerged and rose to high copy-number, entirely replacing the wild-type TE in some populations within a decade (Black et al. 1987; Periquet et al. 1989). These dynamics may reflect selection acting at the level of TEs, with variants outcompeting the ancestral lineage (Le Rouzic and Capy 2006; Robillard et al. 2016; Iwasaki et al. 2020). The clades we identified provide an opportunity to catch such events in progress. We sought therefore to identify TE lineages (using clades as a proxy) that have expanded or contracted in copy number within specific geographic populations, because these might be signatures of selection acting on the TE sequence.

To find clades with population structure we used a Bonferroni-corrected Kruskal-Wallis test to determine which clades rejected the null-hypothesis that their copy number was homogeneously distributed across populations. We found that ∼15% of clades were heterogeneously distributed among the five GDL populations, thus indicating population structure (Figure 4a). Some TE families, such as *Burdock* and *Tart-A*, have few or no clades that are enriched for particular populations, while *Jockey, Copia,* and *Tirant* have many clades with population structure.

**Figure 4.**
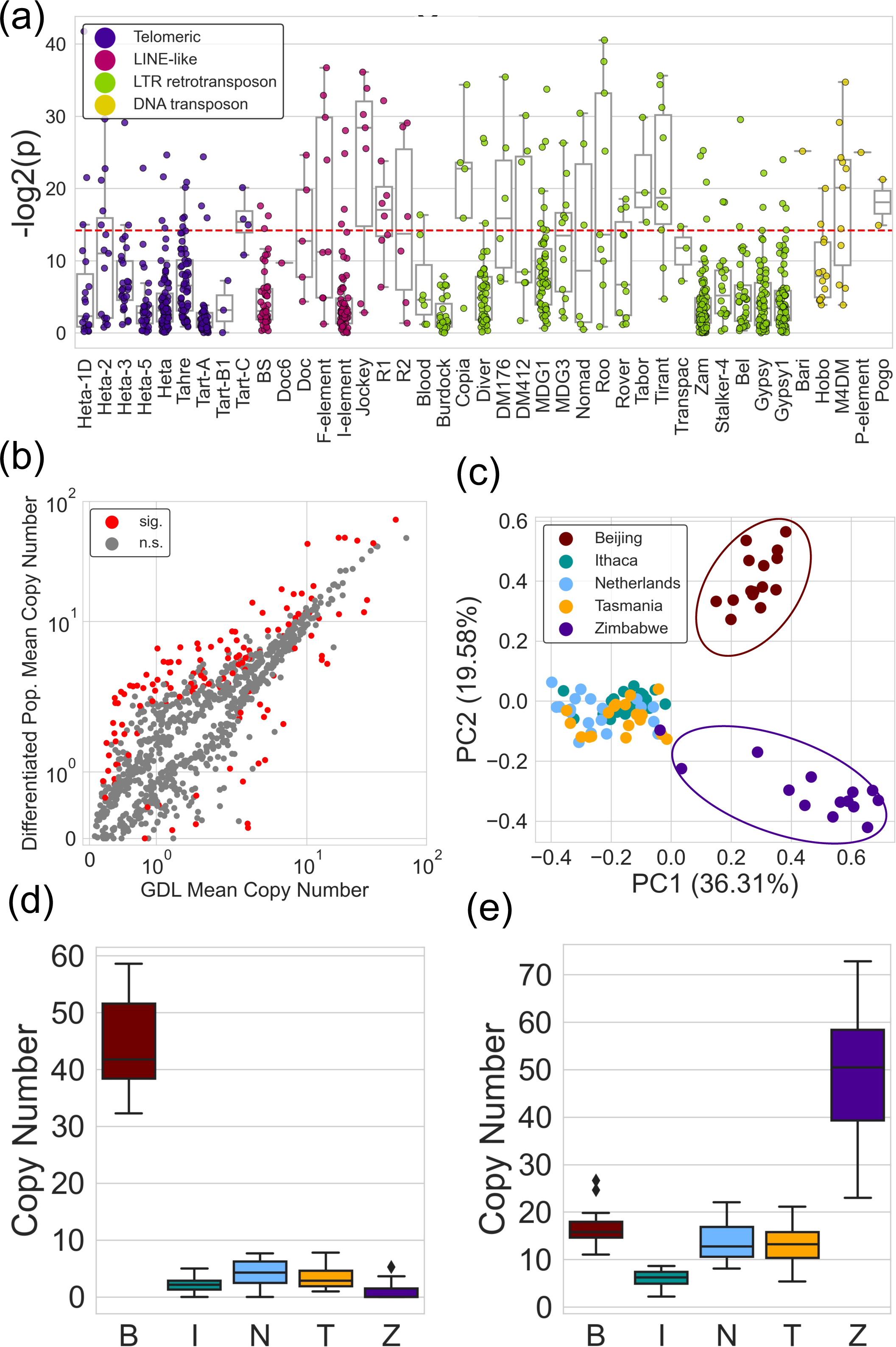
Population structure and variation of TE clades is common. (a) Boxplot showing the result of Kruskal-Wallis tests on the clade copy number between GDL populations for each TE family. Each dot represents the negative log base-2 transformed p-value for a single clade. Red dashed line is the bonferroni corrected critical value. 15% of the clades had a p-value less than the critical value, and showed heterogeneity in copy number between populations 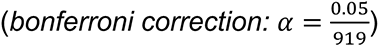. (b) Each dot represents the average copy number of a clade across GDL and the average copy number of the population that is most differentiated from the GDL average. Most differentiated is defined as the greatest absolute difference between the population mean and the GDL mean. Clades are colored by whether they are statistically significant by Kruskal-Wallis test (sig., red), or not (n.s., grey). (c) PCA on the minor allele frequency of *Roo* element SNPs in the GDL. Each dot represents the principal components derived from the minor allele frequencies of an individual. Beijing (red), and Zimbabwe (purple) clusters can be seen. (d) Boxplot of copy number of a *Roo* clade enriched for Beijing (B: Beijing, I: Ithaca, N: Netherlands, T: Tasmania, Z: Zimbabwe). (e) Boxplot of copy number of a different *Roo* clade enriched for Zimbabwe. (B: Beijing, I: Ithaca, N: Netherlands, T: Tasmania, Z: Zimbabwe).

To quantify the extent of the population structure we compared the average clade copy number across the GDL to the average clade copy number of the subpopulation (Beijing, Ithaca, Netherlands, Tasmania, or Zimbabwe) that was the most differentiated from the entire GDL (Figure 4b). Of the clades that were statistically significant by the Kruskal-Wallis test, the most differentiated populations had a clade copy number that was, on average, ∼3.8 copies greater or lesser than the GDL mean. In general, these population structure differences were of modest effect, but *Roo*, *R1,* and *F-element* clades have differences from the GDL mean of ∼10-35 copies. These much larger effect sizes might be driven in part by the very high copy number of these three families throughout the genome.

Our analysis of these summary statistics, although informative, does not reveal in which population(s) a clade is enriched. We therefore employed PCA (Principal Component Analysis) on the matrix of SNP frequencies of each TE family in each individual. This allows us to find which SNPs are driving the population variation within a TE family, as well as to visualize which individuals in the GDL carry similar TE variants. We find strong population structure for *Roo* variants with distinct clusters of Beijing and Zimbabwe individuals (Figure 4c). This population structure is driven by population-specific expansions of clades (Figure 4d, 4e). The Beijing- and Zimbabwe-specific clades are at ∼45 copies (∼33% frequency), and ∼50 copies (∼40% frequency) in Beijing and Zimbabwe, respectively. The Beijing clade is very rare outside of its respective population, ∼1% frequency, which implies that it emerged in East Asia and then expanded in copy number. The Zimbabwe clade, on the other hand, segregates at ∼10% frequency in the other populations, implying a more ancestral origin.

We find an analogous pattern in *Tirant* and *Jockey* variation where there is also strong population structure that is driven by population-specific expansions of clades. Much like *Roo,* Ithaca- and Tasmania-specific clade expansions drive the population structure of *Tirant* variation (Supplemental Figure 5a, 5b, 5c). However, in *Jockey* it is the absence of a clade in Zimbabwe that is found in all other populations, coupled with a Zimbabwe-specific expansion of a different clade, that drives that structure (Supplemental Figure 5d, 5e, 5f). In these three families with notable population structure, it is the presence or absence of a single clade that drives the variation rather than multiple variants expanding within the populations. This pattern could be a reflection of selection favoring the expansion of a single lineage in a population, or simply due to genetic drift. In either case it shows that TE lineages are able to expand in copy number and become endemic in a population, dramatically altering the composition of TE variants within those individuals.

**Figure 5.**
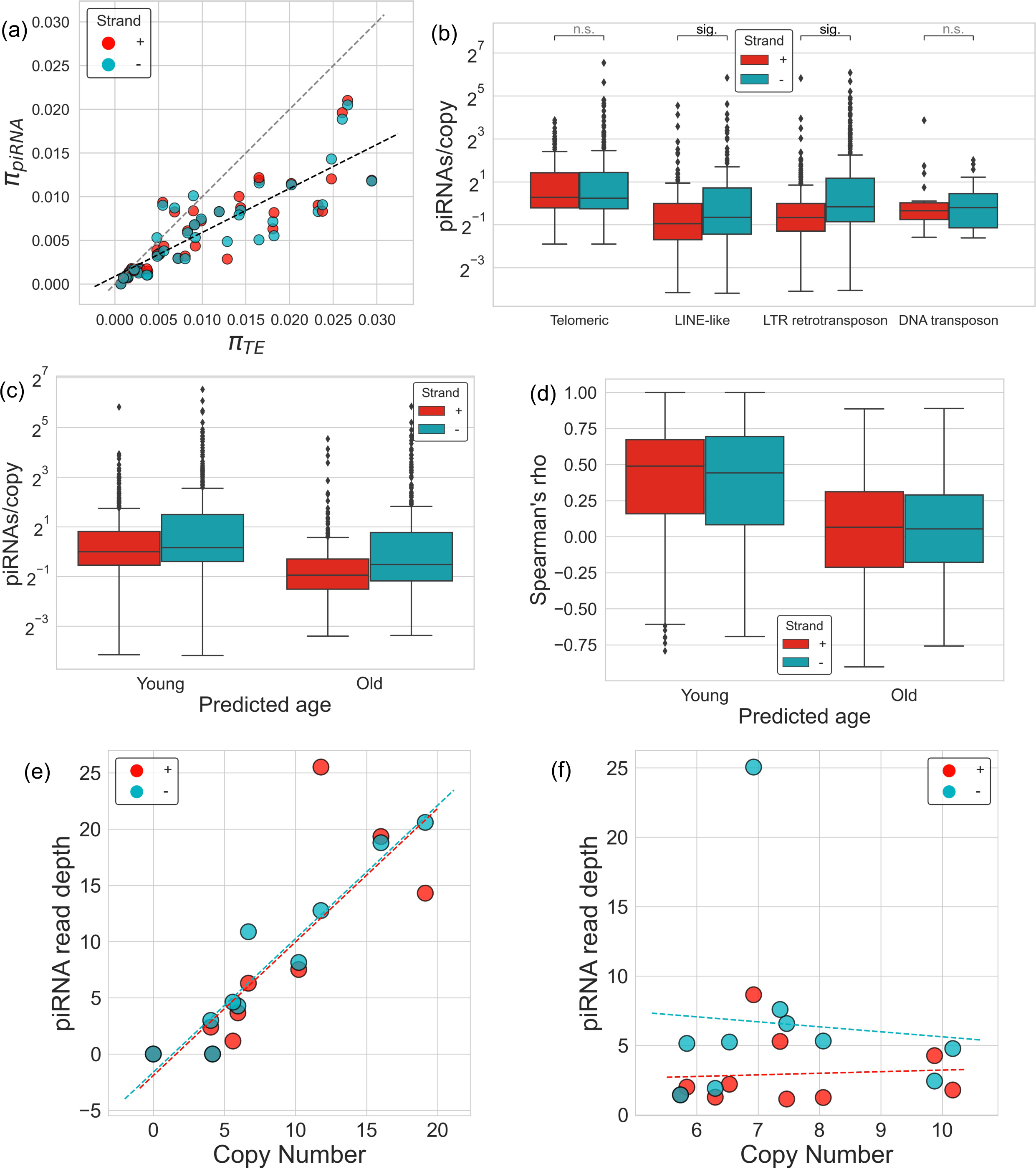
piRNA diversity and the average sense and antisense piRNAs/copy of TE clades for 10 GDL strains. (a) Scatterplot where each point represents the sense (+, red) or antisense (-, blue) piRNA sequence diversity (***π***_piRNA_) for a TE family plotted against the genomic TE sequence diversity (***π***_TE_) of the TE family. The grey dashed line represents the 1:1 expectation of piRNA diversity:genomic diversity and the black dashed line represents a linear fit between the piRNA diversity and genomic diversity. (b) Average clade piRNAs/copy for sense (+, red), and antisense (-, blue) separated by TE class. Significant differences between sense and antisense piRNAs/copy were found in clades for LTR and LINE-like elements (sig.), but not telomeric or DNA transposons (n.s. ; *Wilcoxon signed-rank test*) . (c) piRNAs/copy of putatively young clades (likely active) and old clades (likely inactive). Young clades had greater piRNAs/copy than older clades (+, *Mann-Whitney U p-value* < 0.05; -, *Mann-Whitney U p-value* < 0.05). (d) Spearman’s correlation calculated for copy number and piRNA read depth for putatively young and old clades. Young clades had a greater Spearman’s correlation than inactive clades for sense and antisense piRNA read depth (+, *Mann-Whitney U p-value* < 0.05; - , *Mann-Whitney U p-value* < 0.05). (e) Copy number vs. sense (+, red) and antisense (-, blue) piRNA read depth for a young, recently active *Jockey* clade, and (f) for an old, putatively inactive *I-element* clade.

Not every population-specific expansion of a clade will be as stark as *Roo*, *Jockey*, or *Tirant*. And as we noted above, ∼15% of the several hundred clades discovered show significant heterogeneity in copy number between populations. These clades may not be sufficient to drive variation on a PCA due to their modest effect sizes, but are still significantly different between populations. These small differences may represent stochastic fluctuations in clade copy number between populations, or they may reflect the initial stages of a newly emerging clade in a population rising to high frequency.

### Sense and antisense piRNA pools are diverse and reflect the age of variants

One of the primary mechanisms by which hosts control the proliferation of TEs is through the piRNA pathway. piRNAs are produced in both the sense and antisense direction from two distinct pathways. Antisense piRNAs are generally produced from clusters containing fragments of inactive TEs and target TE transcripts for silencing. Sense piRNAs, in contrast, are derived either from the cleavage of a TE primary transcript, guided by an antisense piRNA, or from dual-stranded piRNA clusters undergoing bi-directional transcription. Generally speaking, antisense piRNAs reflect the potential to silence TE expression, while sense piRNAs reflect the cleavage and silencing of TE transcripts. Sense piRNAs additionally feed back into the production of antisense piRNAs, amplifying the pool of piRNAs targeting that TE sequence (Aravin et al. 2007; Brennecke et al. 2007; Czech et al. 2018). Therefore, the pool of piRNAs a host produces might reflect the genetic variation of active TE families, not just polymorphisms in piRNA clusters.

We asked whether the sequence diversity in the sense and antisense piRNAs (***π***piRNA) tended to be correlated with the sequence diversity of the TEs themselves (***π***_TE_). Sequence diversity of the piRNAs was quantified by aligning ovarian piRNA libraries from 10 strains from the GDL (two from each population) to TE consensus sequences (Luo et al. 2020). For each TE family we pooled together the piRNA reads from the 10 strains to calculate the average piRNA sequence diversity across its consensus sequence and did a similar procedure with the copy-number data. To reduce technical artifacts we only considered SNPs in the piRNA data whose presence was supported by the corresponding genomic data. We found a strong positive relationship between the sequence diversity of a TE family and the sequence diversity in both sense and antisense piRNAs (+: *Spearman’s rho* = 0.90, *p-value* < 0.05; -: *Spearman’s rho* = 0.88, *p-value* < 0.05) (Figure 6a).

The ratio (***π***_piRNA_/***π***_TE_ , dubbed the “piRNA diversity ratio”) in each TE family estimates how well the piRNA diversity reflects genomic diversity. If the ratio is 1, then the piRNAs are as diverse as the genomic loci that they are derived from. A piRNA diversity ratio less than 1 implies that there is greater unevenness in the proportion of variants found in the piRNA pool than in the genomic sequence, such that some variants may be absent from the piRNAs while others dominate. In contrast, a piRNA diversity ratio greater than 1 implies that variants are present in piRNAs at more equal proportions than in the genomic sequence. Across all families of TEs, the piRNA diversity ratio is approximately 0.6 (Figure 5a).

We found that LTR retrotransposons have the lowest piRNA diversity ratio, ∼0.58 for either strand, while DNA transposons and LINE-like elements have ratios of ∼0.7 (Supplemental Figure 6a). A low sense piRNA diversity ratio may reflect TE families with an abundance of old and inactive TE variants, which would produce few sense transcripts and thus few sense piRNAs. Alternatively, these TE variants may be actively transcribing, but not be effectively silenced by piRNAs and therefore few of their transcripts are processed into sense piRNAs. The low sense-piRNA diversity ratio in LTR retrotransposons is surprising given that LTR elements are relatively recent invaders in *D. melanogaster* and therefore many variants of these TE families should be active (Bergman and Bensasson 2007). However, this low ratio is largely driven by older TE families with a high number of old, likely inactive clades (Figure 2c). When we removed LTR retrotransposon families that had old clades, *e.g. Gypsy*, *Gypsy1, Zam*, *etc.,* from the analysis the sense piRNA diversity ratio becomes ∼0.7 (similar to LINE-like elements).

A lower antisense piRNA diversity is likely due to the fact that antisense piRNAs tend to be generated predominantly from piRNA clusters, which contain only a subset of TE variants that are not necessarily representative of all TE variation. *P-element* has the lowest piRNA diversity ratio of all families, ∼0.01, likely reflecting the low genetic diversity of *P-element* (***π*** = ∼0.00076) and their recent invasion in *D. melanogaster* (Kidwell 1983). Few *P-element* variants are segregating in the population and even fewer presumably have been captured by piRNA clusters. Differences in the piRNA diversity ratio between classes may also be reflective of differences in the rate of mutation. High mutation rates during transposition could increase ***π***_TE_, but not increase ***π***_piRNA_ if the piRNAs are derived from a small subset of insertions, such as piRNA clusters.

Although most TE families had a low piRNA diversity ratio, *R2, Blood* and *Hobo* had a piRNA diversity ratio > 1 for one or both strands, and *Roo* was about 1.5x for both strands. These results were not due to an abnormally low average genomic diversity (∼0.005 - 0.01). A high sense piRNA diversity ratio implies that many TE variants are transcribed and targeted by piRNAs, while the high diversity of the antisense piRNAs indicates that most variants are present in piRNA clusters or are producing de novo antisense piRNAs.

Given that piRNA content was generally less diverse than the TEs themselves, we wanted to determine which variants were contributing to the diversity of the piRNA pools. Therefore, we calculated the number of mapping sense and antisense piRNAs that contained the lineage-informative SNPs of a clade per copy of that clade and averaged this ratio across the 10 GDL strains. A clade that is both being regulated by piRNAs and being used to produce piRNAs should have high quantities of both sense and antisense piRNAs.

With the exception of the telomeric TEs, antisense piRNA read depth of lineage-informative SNPs was more abundant than sense. This difference was most stark in LTR retrotransposons where over 4x more antisense than sense piRNA reads/copy contained lineage-informative SNPs (Figure 5b, *Wilcoxon signed-rank test: p-value* < 0.05). This suggests that many of these clades have been incorporated into piRNA clusters and are producing antisense piRNAs. The telomeric TEs not only showed similar proportions of sense and antisense piRNAs per copy, but also generally had more piRNAs per clade copy than other TEs, likely reflecting the fact that piRNAs targeting telomeric TEs are generated from the telomeric TEs themselves rather than distinct piRNA cluster loci (Radion et al. 2018). Newly evolved telomeric TE variants do not have to insert by chance into an existing piRNA cluster or become converted into a *de novo* piRNA cluster, but instead can be immediately incorporated into the pools of antisense piRNAs.

We find that many clades produced few or no piRNAs, but there were some, such as in *HeT-A5*, that produced over a hundred piRNAs/copy (Supplemental Figure 6b). The low piRNA diversity ratio we observed for TEs such as *HeT-A5* may therefore reflect the inclusion of only a subset of clades in the primary or secondary piRNA pathway, and this inclusion may not be representative of the copy number of those clades. It is clear that some clades are more likely to be present in the piRNA pool than others. We therefore hypothesized that recently active variants might be more readily targeted by host piRNAs and therefore produce more piRNAs/copy. We used the above described classifications of young and old clades based on the Poisson fit of their copy-number distributions and analyzed the piRNA abundance of the two groups. We found that young clades have significantly higher sense piRNAs/copy, fitting our prediction that these clades are indeed actively transcribed and therefore likely transpositionally active (*Mann-Whitney U: p-value* < 0.05) (Figure 5c). This also held for antisense piRNA read depth, although the difference was less pronounced (*Mann-Whitney U: p-value* < 0.05). It is clear that although young, recently active clades are more readily used as a substrate by the primary piRNA pathway, there are still many older clades that generate antisense piRNAs, perhaps representing old heterochromatic piRNA clusters containing inactive variants. This finding is consistent with previous analyses that showed older TE families produced less piRNAs than canonical younger families (Kelleher and Barbash 2013). We note that our classifications using the theoretical expectations of the copy-number distribution are imperfect, and some “older” variants may still retain transpositional activity despite their age.

Previous models of evolutionary arms races between TEs and piRNAs predict a positive correlation between the copy number of invading elements and the production of antisense piRNAs, because there is selection on the host genome to silence these elements (Luo et al. 2020). Therefore, for each clade we calculated the Spearman’s rank correlation coefficient between its copy number and piRNA read depth for the 10 GDL strains, and compared these values between active and inactive clades. We found that young clades were significantly enriched for positive correlations in both sense and antisense piRNAs (Figure 5d, 5e, 5f; +: *Mann-Whitney U*: *p-value* < 0.05; -: *Mann-Whitney U*: *p-value* < 0.05). We also found that there were 61 and 58 young clades that had a statistically significant correlation between copy number and antisense and sense piRNA read depth, respectively, while only 2 and 4 old clades were statistically significant (*Benjamini-Hochberg: FDR = 10%*). Of the young clades many belonged to telomeric TEs, or recently active LTR and LINE-like retrotransposons, like *Jockey*, and *Tirant*.

Overall, our analysis of piRNA sequence variation shows that host piRNA content changes to respond to the emergence of variant TEs, and that not all variants are represented in the piRNA pool. Young, putatively active TE variants are disproportionately represented in the sense and antisense piRNAs, suggesting that host genomes may be responding to the evolution of new TE lineages.

## Discussion

### Clade inference provides a new tool for understanding the evolution of TEs

We have developed a technique for inferring clades within TE families by leveraging population genomics datasets and heuristic statistical methods. This approach bridges a significant gap in the field of population genomics by obtaining information about TE family substructure from existing short-read datasets. Simulations show that this method reliably identifies clade structures that are consistent with the TE genealogy under a wide parameter space, and we also validated TE clade inferences in *D. melanogaster* by checking them against PacBio genomes. We then used the clade designations from data on natural *D. melanogaster* populations to infer aspects of TE dynamics and host responses.

The clade calls should be interpreted with some caveats in mind, however. The clusters of lineage-informative SNPs are markers that distinguish clades in the TE genealogy, not the complete set of SNPs within a full-length insertion. Given that two clades may descend from the same ancestral lineage, clusters of SNPs may co-vary within insertions but still be called as distinct TE clades. This behavior reflects a trade-off between merging versus splitting clades and depends on the chosen false-positive rate. This choice will affect the number of clades called for a given TE, but does not likely change the relative proportions of clades among different TEs, which our simulations show are robust to perturbations in clustering parameters. The technique we have developed can be readily applied to any organisms where population-level short-read genomic sequence data and libraries of TE consensus sequences exist.

### Extensive population variation of TE clades

The study of the genetic variation of TEs has previously been largely relegated to reference genome assemblies. By applying population genetic theory to the copy-number distribution of clades, we found that a majority of clades (56%) were young and recently active. This is not wholly unexpected as most of the TE families we assayed have sequence diversity and population insertion-site frequency spectra that reflect invasion, activity and/or selection against TE insertions that are recent (Kofler et al. 2012; Kelleher and Barbash 2013; Kofler et al. 2015). However, the classifications of young and old clades do not necessarily reflect the true transpositional activity of those elements, and although we orthogonally validated our inferences of age (Supplemental Figure 4), the effects of demography, population structure and TE life history complicate our inferences and may give rise to errors. Ultimately, our inference of age is based on the expected association of reduced transpositional activity and a broad insertion-site frequency spectrum, which will not always hold.

Despite this, our analysis found that some putatively young clades, such as in *Jockey* and *DM412*, have expanded in copy number dramatically across all populations, accounting for ∼40% of all insertions. Other young clades have expanded only within a subset of the populations, sometimes to 3-4x higher copy number than other populations. Interestingly, Tasmanian-specific SNPs for *Tirant* have been previously observed, but our study is the first to put this observation in the context of the emergence and expansion of a TE lineage (Schwarz et al. 2020).

This begs the question: what drives these differences in copy number? One possible cause of local copy-number expansions is the acquisition of adaptive polymorphisms that increase transposition rate. For example, *hobo* elements in *D. melanogaster* with 5 copies of an internal repeat are less active than variants with 3 copies (Souames et al. 2003). There are also segregating polymorphisms in the human *LINE-1* element that account for ∼16 fold differences in transposition rate (Lutz et al. 2003; Seleme et al. 2006). Polymorphisms could affect the transposition rate by increasing efficiency during replication or by evading host genome- silencing mechanisms (Han and Boeke 2004; Cosby et al. 2019). A more transpositionally efficient variant would eventually displace other variants as it increased in copy number within that population (Le Rouzic and Capy 2006).

It is also possible that differences in clade copy number between populations are caused by neutral processes. Founder effects and geographic isolation could affect the copy number and composition of variants within a population, thus creating population structure (Jurka et al. 2011; Lerat et al. 2019). Population genetic simulations of TEs competing within a population provide a future way to explore these hypotheses.

### Antisense piRNA production of variants may be adaptive

Although the piRNA system can quickly respond to the invasion of TE families into naive populations by producing antisense piRNAs specific to those new invaders (Kofler et al. 2018), its ability to change in response to the emergence of new variants of a TE family has been underexplored. We have shown that the piRNA defense system is surprisingly malleable and seems to often respond to the emergence of new variants by incorporating those variants into antisense piRNAs. The presence of a variant in the antisense piRNAs indicates inclusion of that variant in a piRNA cluster, and may reflect the propensity for the host to silence those variants. We found that, in general, antisense piRNAs had less sequence diversity than genomic TE insertions and that young, recently active clades were overrepresented in the antisense piRNAs. This is consistent with previous findings that showed a bias for piRNA silencing of active human *LINE-1* elements (Lukic and Chen 2011). In *D. melanogaster*, a positive relationship was found previously between indicators of transposition activity for TE families and their antisense piRNA abundance. However, this relationship seemed to be driven by the removal of inactive TE families from the piRNA pool rather than an increase in the silencing of active elements (Kelleher and Barbash 2013). Our analyses largely concur, as older clades of TEs are significantly less represented in the piRNA pool than younger clades.

Furthermore, the malleability of piRNA content might be beneficial to the host as positive correlations were found between the copy number of young clades and their antisense piRNA read depth. Such positive correlations are predicted under an evolutionary arms race model with strong piRNA silencing (Luo et al. 2020). Under this arms race model, the copy-number expansion of TEs is counteracted by a corresponding expansion of piRNA clusters that are capable of silencing those elements. Under this model the selective effect of TE insertions must be deleterious and the efficiency in silencing must be high (Luo et al. 2020). Recent analyses of TE family copy number in *D. melanogaster* laboratory and natural populations found positive correlations between piRNA read depth and copy number in 6 out 105 families analyzed. These were mostly young and recently expanding TE families, including *P-element* and a handful of telomeric TEs (Luo et al. 2020; Saint-Leandre et al. 2020). By considering the copy-number variation of clades within TE families, our analyses provide much wider evidence of the expected correlation, with antisense piRNA production correlated with copy number for 61 TE clades in 22 out of the 41 likely active TE families considered. These include many telomeric TE clades, for which the positive correlation may have a distinct mechanistic explanation: as nearly all telomeric TEs are found at the telomere ends and do not insert at pericentromeric piRNA clusters, the piRNAs must be generated from the telomeres (Radion et al. 2018). However, we discovered that other active TE families, including *Roo, Jockey, R1,* and *Tirant*, also show this correlation and were not detected previously. The increased power in our analysis would be expected if active lineages preferentially display this correlation between piRNA read depth and TE copy number, with family-level analyses losing statistical power due to the aggregation of young and old clades. This highlights the importance of integrating sequence polymorphisms into the analysis of TEs and the utility of our clade inference method.

Because the arms-race model predicts significant positive correlations between antisense piRNA abundance and copy number when the strength and efficiency of piRNA silencing is high, it is possible that the host produces antisense piRNAs that are specific to recently active clades to increase silencing efficiency (Luo et al. 2020). Those piRNAs that have perfect sequence complementarity to their targets might have higher specificity in binding and therefore increased silencing efficiency. In *C. elegans* and *D. melanogaster,* deletions or polymorphisms in piRNA binding sites on a transcript can reduce or eliminate silencing if sufficiently diverged from the piRNA sequence (Post et al. 2014; Zhang et al. 2018; Kotov et al. 2019).

Given that piRNA silencing efficiency is affected by sequence complementarity, significant enrichment for recently active TE clades in antisense piRNAs may be driven by natural selection acting to increase the frequency of piRNA clusters segregating in the population that contain active TE variants. piRNA clusters can be highly polymorphic in TE content, rapidly turnover in sequence and be enriched for young, recently active TE families (Assis and Kondrashov 2009; Zanni et al. 2013; Wierzbicki et al. 2020; Zhang et al. 2020; Gebert et al. 2021 Jul 30). Many distinct piRNA clusters are therefore likely to be segregating in *D. melanogaster* populations, each with distinct compositions of TE families and variants. The piRNA clusters that contain newly emerging variants may be selected for if they more efficiently silence novel variants, thus increasing in frequency. Although there is strong selection to maintain piRNA mediated silencing of TEs in a population (Bergthorsson et al. 2020), the effective strength of selection on piRNA clusters containing new variants is unclear. Full de-repression of TEs is only seen when they are > ∼10% diverged from their respective piRNA sequences (Kotov et al. 2019), and the clades that we inferred are only diverged from the consensus sequence at a handful of sites. But it is conceivable that modest divergences in sequence could create an unequal regime of silencing between variant TEs.

Alternatively, enrichment of active TE variants in the piRNA pool may be independent of selection. Instead, transcriptional activity of euchromatic TE insertions may drive variation in piRNA pools. This epigenetic model is plausible because the transgenerational inheritance of piRNA cluster expression is dependent on maternally deposited piRNAs that trigger the production of “secondary” piRNAs from euchromatic TE insertions that are thought to then feed back into germline piRNA clusters (Brennecke et al. 2008; Le Thomas et al. 2014; Senti et al. 2015). Furthermore, TE insertions targeted by piRNAs may be converted into *de novo* piRNA clusters capable of producing sense and antisense piRNAs through recruitment of the *Rhino-Deadlock-Cutoff* (RDC) complex, a process called licensing (Olovnikov et al. 2013; Mohn et al. 2014; Shpiz et al. 2014). Euchromatic insertions of young TE variants will more likely be transcriptionally active than insertions of older variants, making them a more prominent target of silencing by maternal or germline antisense piRNAs and therefore increasing the likelihood of RDC recruitment. This licensing mechanism could “switch on” the activity of piRNA clusters that contain active TE variants, thus establishing a transgenerational change in piRNA content without altering the frequencies of piRNA clusters in the population. In fact, a repetitive locus can be experimentally converted into a heritable piRNA cluster in certain environmental conditions (de Vanssay et al. 2012; Casier et al. 2019). These *de novo* germline piRNA clusters will then produce piRNAs that could be maternally deposited to the next generation and reinforce piRNA cluster identity. The formation of *de novo* piRNA clusters as a primary mechanism by which piRNA pools change in populations over time is a favorable hypothesis, as canonical germline piRNA clusters can be dispensable in the silencing of certain TE families (Gebert et al. 2021 Jul 30) and young clades tend to be found less often in heterochromatin than old clades, making their capture in canonical heterochromatic piRNA clusters less likely (Supplemental Figure 4).

In the first model, selection plays a major role in determining the piRNA content in a population and the enrichment of variant-containing antisense piRNAs is strictly adaptive. But in the epigenetic model, the enrichment is not necessarily adaptive. The piRNA content may shift to bias more active variants due to variation in piRNA cluster activity through the formation of *de novo* clusters, but these variant-containing piRNAs need not be more efficient at silencing TE variants. It is possible that this epigenetic variation is beneficial for the host, or it may be a byproduct of the mechanisms by which piRNA clusters are inherited. These two models are not mutually exclusive, and the underlying observations reveal a fundamental aspect of the TE-host relationship, whereby a new TE variant emerges in a population, increases in copy number, and then is used as a substrate by the host genome to produce novel antisense piRNAs.

## Methods

### Aligning short-read data to TE consensus sequence using ConTExt and estimating copy number

85 short-read libraries from the Global Diversity Lines were aligned to a curated index of RepBase TE consensus sequences and the *D. melanogaster* release 6 reference genome (Bao et al. 2015; Grenier et al. 2015; Hoskins et al. 2015; McGurk et al. 2020 Dec 22), using *ConTExt* following the parameters in (McGurk and Barbash 2018). From this output we estimated the copy number of each position for every TE consensus sequence from the read depth, as described in (McGurk et al. 2020 Dec 22), and used the read pile-ups to calculate allele frequencies. Copy-number estimates and allele frequencies were generated for each of the 85 short-read libraries for each TE consensus sequence. For Long Terminal Repeats (LTRs), or Perfect Near-Terminal Repeats (PNTRs) sequences the consensus sequence for the repeat unit is too short for copy-number estimation from read depth, so in these cases we used the median copy number from the internal sequence as the estimate of copy number. Additionally, we appended the LTR/PNTR copy number, and allele frequency data to the end of the internal sequence in order to be able to infer SNPs on the LTR/PNTR that co-occur with internal SNPs.

### Filtering reads by mapping quality

When creating the copy number and allele frequency matrices, reads aligned to the TE consensus sequences were filtered for mapping quality as described in (McGurk et al. 2020 Dec 22). In brief, rather than using the Bowtie2 mapping quality scores we derived our own metric of filtering ambiguous reads based on the percent identity of the read to the primary (AS) and secondary (XS) alignments. We chose to filter reads in this way because we expect that many reads will be diverged from the consensus sequence if they are derived from polymorphic elements, and we would like to retain that information. We first convert the alignment score of the read alignments to the percent identity to the consensus sequence by assuming all penalties are due to mismatches, and then use these percent identities for AS and XS to compute a score:

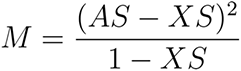

Which reflects the distance between the primary (AS) and secondary (XS) alignment penalized by the divergence of AS to the consensus sequence. If secondary alignments are reported by Bowtie2 we require this score to be greater than 0.05 for the alignment to be included in the analysis. If only a primary alignment is found the alignment must be less than 20% diverged from the consensus sequence to be included.

### Calculating sequence diversity of TE families using NGS data

From the copy number and allele frequency data derived from the read alignments to the TE consensus sequence we calculated the sequence diversity at each position in the alignment. We multiplied the copy-number matrices by the allele frequency data to generate the estimated number of copies for all alleles across the sequence of the TE consensus sequence, and removed alleles with a copy number < 0.5 as we assumed these low values reflected sequencing errors. We calculated the allele copy number of a TE family for each strain’s alignments, as well as pooling allele copy number for all strains belonging to the same population (*e.g.* all Beijing strains), and pooling all strains to obtain global allele copy-number data. We next estimated sequence diversity at each position for each strain, population, and the entire dataset using the allele copy-number data as:

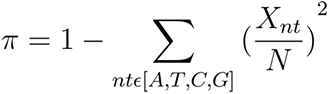

Where *N* is the total copy number at that position and *X_nt_* is the copy number of an allele. When calculating the sequence diversity of the piRNA reads we performed the same procedure, but used a matrix of read counts rather than copy number and did not include any alleles with a copy number < 0.5.

### Inferring TE Clades

We developed a method to infer the co-occurrence of SNPs within a TE sequence by finding positive correlations in copy numbers between SNPs across multiple individuals. We performed this inference on a set of 41 recently active TEs (Supplemental File 1, https://github.com/is-the-biologist/TE_CladeInference). For this approach we only included positions within a TE that had a within-population sequence diversity > 0.1; or had an overall sequence diversity > 0.1. This would be equivalent to filtering out positions where the major allele is present in greater than 95% of copies. After initial diversity filtering, we obtained the copy number estimates of each allele by taking the proportion of reads that mapped to each allele and multiplying it by the estimated copy number at that position. For each position we determined the major allele as being the allele with highest copy number across the entire dataset and then extracted the copy number of the three minor alleles for every strain at every position. The result of this is an *S* x *N* matrix, where *N* is the number of minor alleles that passed our diversity criteria and *S* is the number of strains in the dataset. Each element of this matrix contains the copy number estimates for that allele for each strain. To reduce the rate of false positive correlations caused by low-copy-number alleles, we required that an allele must be present in at least 10 strains to be considered. Additionally, because we are only interested in high frequency alleles, we required alleles to be present in at least 10% frequency either across the GDL, or within a population.

Additionally, we removed strains from the *S* x *N* matrix that were determined to be outliers in copy number as determined from (McGurk et al. 2020 Dec 22). These outliers do not represent the natural variation in copy number and instead represent TE copy-number expansions that likely occurred during the inbreeding process of the strain. These massive expansions break assumptions of our method by allowing situations where distinct TE subfamilies may co-expand in copy number and correlate while not existing on the same TE sequence. We perform this data processing for each active TE of interest.

To identify lineage-informative SNPs we perform Hierarchical Clustering with average linkage using a correlation distance on the *S* x *N* matrix (Using the R package *pheatmap*). This clusters together alleles that correlate in copy number. We use this to both seriate a correlation matrix of the alleles, and to directly call clusters by cutting the dendrogram at a correlation distance optimized for each TE, as described in *Choosing a Distance Cut Off for Hierarchical Clustering.* Clusters of minor alleles with more than one allele are lineage-informative SNPs that can be used to distinguish TE clades.

### Choosing a Distance Cut Off for Hierarchical Clustering

We validated a critical assumption of our inference model: that SNPs that are physically linked within the same TE sequence will co-vary in copy number, while SNPs that are not physically linked on unrelated TE sequences will not. To do this we asked whether the SNPs in different TE families are positively correlated, expecting that because the SNPs on different TEs are physically unlinked there will be little to no positive correlation between them. We computed the correlation for every pairwise combination of active TEs in our dataset and found, as expected, the correlation of SNPs taken from the same TE family (average Pearson’s *r* = 0.23) are generally much greater than from unrelated families (average Pearson’s *r* = 0.02) (Supplemental Figure 7a). There is an elevated number of positive correlations between SNPs on different telomeric TEs (average Pearson’s *r* = 0.11), likely due to them being linked together in large multimeric arrays exclusively at the ends of chromosomes. There are also very strong positive correlations of SNPs in the *Bari* element (Pearson’s *r* = 0.69) and *P-element* (Pearson’s *r* = 0.68) that are driven by the small number of SNPs segregating in those two families. There are only three SNPs that passed filtering for *Bari* and four for *P-element*, which represent one clade in each TE family.

To justify the correlation cut-off criteria in our Hierarchical Clustering, we generated a null distribution of correlations by permuting the order of the rows of each column of the filtered sets of minor alleles and calculating the pairwise correlation of these permuted SNPs. We performed this operation 1,000 times for each TE. We calculated the pairwise correlations of the un-permuted filtered sets of minor alleles and denoted this as the Test distribution. Due to the large number of pairwise comparisons performed in the clustering (869,042 pairwise correlations) we sought to correct the false positive rate by performing Bonferroni correction on our critical value ( ) by dividing by the number of pairwise comparisons. This critical value is subtracted from 100 to obtain a percentile that we use to determine cutoffs in the hierarchical clustering, a so-called “Critical Percentile”. We pooled the Test and Null distributions across all TEs of interest and computed the correlation value at the “Critical Percentile” in the Null distribution, *r* = 0.59 (Supplemental Figure 7b). Although stringent, we found that 6.38% of the SNPs in the Test distribution had a *r* > 0.59.

We further sought to justify our cut-off by examining the individual Null and Test distributions for each TE and comparing the Null distributions between elements (Supplemental Figure 7c). We observed that there was some degree of variability of the Test and Null distributions for each of the TEs. The “Critical Percentile” of the Null distributions fell between a correlation value of ∼0.43 - 0.93 across the samples. Therefore, we optimized the hierarchical clustering distance cutoff for each TE by setting it to the correlation at the “Critical Percentile” for each Null distribution.

### Downsampling GDL NGS data

In order to benchmark our method’s ability to infer clades under different sample sizes, we downsampled the number of GDL strain libraries and compared the resulting clades to the clades inferred using the full 85 GDL strain libraries. We downsampled strains to 75, 50, 25, 15, 10 and 5 by randomly removing strains from the post-filtering allele copy number matrices and inferred clades in all 41 TEs using the same set of correlation cut-offs used for the original inferences with all 85 strains. We then calculated the Rand Index of the downsampled inferences considering the original inference using 85 strains to be the “true” set and the downsampled inferences as the “test” set (Supplemental Figure 3a). The Rand Index (*RI*) may be viewed as the proportion of clusterings in the “test” set that were found in the “true” set, or:

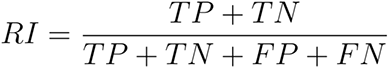

Where, TP is true positives, TN is true negatives, FP is false positives and FN is false negatives.

Overall, RI was robust to sampling size for many TE families (RI between ∼0.8-0.9), but decreased once sample sizes dipped below 50 strains. In LINE-like retrotransposons and telomeric TEs the effect of sample sizes was much stronger, likely due to the fragmented nature of the TEs breaking down positive correlations in the data (See Simulations of TE copy-number data to benchmark clade inference for details). We also found that the total number of clades inferred using the downsampled data varies considerably (Supplemental Figure 3b). The accuracy of clustering is limited when the number of strains drops below 50 and caution should be taken in interpreting the output from this method under those circumstances, especially when inferring clades in LINE-like retrotransposons.

### PacBio Data and Alignment of TE Consensus Sequences

For analysis of PacBio data we used 13 DSPR founders, the Oregon-R PacBio genome, and five PacBio genomes from the GDL (Long et al. 2018; Chakraborty et al. 2019). RepBase consensus sequences of 41 TEs of interest were processed by substituting ambiguous base calls with a non-ambiguous nucleotide, and then aligned to the PacBio genomes using BLASTn (Altschul et al. 1990; Altschul et al. 1997; Camacho et al. 2009; Bao et al. 2015). For retrotransposons with LTRs or PNTRs we aligned only the internal sequence to simplify the amount of downstream processing of the alignments, and because the majority of SNPs reside in the internal sequence. Alignments were output as XMLs to be analyzed downstream. We also extracted the sequences of each of the alignments as fasta files to be used to construct phylogenies (Supplemental File 2, https://github.com/is-the-biologist/TE_CladeInference).

### Constructing TE Phylogenies from PacBio Data

We constructed phylogenies of TEs by using TE fasta sequences extracted from PacBio genomes, and then annotated the tips of the phylogeny with inferred clades from the GDL short-read data. We constructed phylogenies for all 41 TEs analyzed (Supplemental File 3, https://github.com/is-the-biologist/TE_CladeInference). We first extracted fasta sequences of each insertion in the PacBio genomes by taking the sequences from the alignments described above. We excluded TE sequences that were less than 75% full length and then generated a multiple sequence alignment of the remaining sequences using clustalOmega (Sievers et al. 2011). A phylogeny of the sequences was constructed using maximum likelihood and model fitting with the tool iqTree2. A consensus tree was built using 1,000 bootstrap replicates (iqtree2 -s {input} -bb 1000) (Kalyaanamoorthy et al. 2017; Hoang et al. 2018; Minh et al. 2020). We used this consensus tree to generate cladograms, and colored tips of the phylogeny by which clades they belonged to. We considered sequences to belong to a clade if they contained two or more lineage-informative SNPs (Supplemental File 3, https://github.com/is-the-biologist/TE_CladeInference).

### Effect of Clustering Parameters on Phylogenetic Distance of Clades

We sought to understand how inferred clades relate to the phylogeny of TE sequences. In particular, we were interested in determining the effect of the correlation cut-off used for clustering on the phylogenetic distance between insertions belonging to inferred clades. We expected that more stringent correlation cut-offs would produce clades with closely related sets of insertions, while lenient cut-offs would produce clades with more distantly related insertions.

To determine if this was true we inferred clades in real and simulated data, varying the correlation cut-off from lenient (Pearson’s *r* = 0.1) to stringent (Pearson’s *r* = 0.9), and for each clade inferred we summed all pairwise cophenetic distances of each tip (TE insertion). Tips belonged to a clade if their sequence contained 2 or more lineage-informative SNPs of that clade. The calculated distance was normalized by dividing by the sum of all pairwise cophenetic distances between tips in the tree, producing a value between 0 and 1. A value near 0 means that tips belonging to that clade were closely related and composed a relatively small proportion of the phylogenetic tree, while a value near 1 means that the tips were distantly related and composed a large proportion of the tree. We normalized the phylogenetic distance to make comparisons between TE families easier and to be able to interpret phylogenetic distances as a proportion of the total branch lengths in a given tree.

We report the clade-wise average of the normalized distances for each TE family under correlation cut-offs between 0.1 and 0.9 (Supplemental Figure 1a). Overall we found that our expectation holds true, with insertions belonging to clades inferred under relaxed clustering cut- offs being more distantly related than those inferred under stringent cut-offs. Relaxed cut-offs will generate clades composed of many SNPs, collapsing much of the phylogeny into one clade, while stringent cut-offs produce clades with SNPs that are more strongly correlated and closely related. This can be seen clearly in the inferred clades of simulated data (Error rate = 0.1%; Full length elements), where we show the normalized distance of each clade under the regime of correlation cut-offs (Supplemental Figure 1b). We additionally show the simulated phylogeny annotated with clades inferred at Pearson’s *r = 0.6* (Supplemental Figure 1c).

From this analysis we observe that there is a relationship between how closely related insertions bearing two SNPs are and the degree of correlation between the copy-number of those SNPs. A few TE families are exceptions, with insertions in a clade being distantly related but having SNPs that are very strongly correlated (Supplemental Figure 1a). This is illustrated most clearly in the *Jockey* element, where lineage-informative SNPs composing the clade “Cluster_7” are strongly correlated (Pearson’s r > 0.9), but insertions belonging to this clade are widely distributed along the phylogeny (Supplemental Figure 1d). Almost every *Jockey* sequence used to build the phylogeny belongs to “Cluster_7” hence its wide distribution. We show the cladogram of *Jockey* annotated with clade calls from the correlation cut-off used for the downstream analysis of GDL data in this manuscript (Pearson’s r = ∼0.55) to concretely visualize the phylogenetic distribution of clades (Supplemental Figure 1e).

### Validation of GDL clades using PacBio genomes

We used the alignments of TE consensus sequences to the PacBio genomes to recover our clades inferred from the GDL short-read data. To do this we first needed to extract phased haplotypes from our alignments. We took the alignments and recorded the position on the consensus sequence and the nucleotide at each position found in each TE insertion of the PacBio genomes. We accounted for gaps created by insertions and deletions by correcting the position, or adding missing values, respectively. The result is a sequence for each alignment that records the nucleotide at each position of each TE insertion found in the PacBio genomes relative to the position in the RepBase consensus sequence.

We then checked for the presence of lineage-informative SNPs discovered from the GDL short-read data in all TE insertions in the PacBio genomes by querying those SNPs against the aforementioned alignments for a given TE family. For each clade we removed aligned PacBio sequences if they had a deletion at one of the positions of a lineage-informative SNP. On occasions when a SNP was not found in any aligned sequence, that SNP was removed from the analysis. Additionally, we removed clades from the validation if less than two of the SNPs were detected in the PacBio data. A total of 1,719 out of 4,383 SNPs were removed from the PacBio analysis. The proportion varied among TEs, with some elements like *Tart-A* or *P-element* having 95-100% of SNPs missing while other elements like *Doc* had no SNPs missing. The SNPs absent in the PacBio genomes may be present in TE insertions in heterochromatin and were thus unassembled, or may not be present in the 19 strains sequenced using PacBio. It should also be noted that for retrotransposons only their internal sequence was aligned to the PacBio genome and so SNPs that reside on the LTR/PNTR were not used in this verification.

We recorded the frequency at which each of the filtered clades were found in the PacBio genomes and found that approximately 70% of the clades inferred from the GDL short-reads were found within the PacBio alignments (Figure 2a). We also performed the same analysis including clades where no SNPs were detected in the PacBio genomes and found that while the total percentage of clades detected decreases to 38%, the trends on the percent clades validated for each TE are similar (Supplemental Figure 8a). Missing clades not found in the PacBio genomes may reflect several distinct technical and biological issues. Firstly, some sets of clades do not have perfect linkage between all of their SNPs. This can occur when related lineages that share SNPs are segregating within the population. The clustering algorithm is unable to distinguish these multiple lineages and clusters them together, because they share a significant portion of their SNPs. The other possibility is that the lineage, or a subset of SNPs in the lineage, are rare or specific to a population in the GDL, and were not sampled in the PacBio genomes.

To more finely describe the co-occurrence of SNPs in clades, we computed the pairwise Jaccard Score of SNPs within and between inferred clades. The Jaccard Score is computed as the number of times two SNPs occur together in a TE insertion in the PacBio genomes divided by the total number of times that either one or both SNPS are present. We converted these scores into a Jaccard Distance by simply calculating 1 - Jaccard Score, such that two SNPs that always co-occur will have a distance of 0 while two that never co-occur have a distance of 1.

We used these Jaccard Distance matrices to quantify the cohesiveness and separation of lineage-informative SNPs by computing Silhouette Scores. Silhouette Scores are a common metric for evaluation clustering performance and are calculated as being the mean distance between an individual in a cluster and its other cluster members subtracted by the mean distance between this individual and the members of the closest neighboring cluster. These values are then normalized such that they are bounded between -1 and 1, where positive scores imply that SNPs within a cluster, or clade, co-occur with each other more often than they co-occur with SNPs from a neighboring clade, and negative scores imply the opposite.

To explore how well the identified clusters reflect true clades of insertions we classified clades into four categories by comparing the Silhouette Score of clades to their frequency in the PacBio data (Supplemental Figure 8b). “Full clade” is analogous to a true positive. These clades have positive Silhouette Scores at a frequency greater than 0, *i.e.* clustering quality is good, and this arrangement of SNPs is found in the validation data. “Multiple derived lineages” also have a positive Silhouette Score, but their frequency is 0. Their interpretation is complex with no easy analogy, however we reason that the sets of SNPs co-occur in multiple lineages and the algorithm may be merging these multiple lineages into one cluster. “Incomplete clade” is similar to a false-negative, or under-clustering. In this instance, the Silhouette score is negative, but the frequency is greater than 0. These arrangements of SNPs are found to exist in the validation data, but may be a result of splitting lineages, or can reflect a high degree of relatedness with other clusters. “Errors” are likely false-positives. They have a Silhouette Score that is negative and a frequency of 0. In these cases the SNPs are found in the validation set independently, but do not co-occur in the same TE insertions together, although some subset of the SNPs may co-occur.

Using these classifications we found that 41.4% of inferred clades are “Full clades”, 5.5% are “Multiple derived lineages”, 28.4% are “Incomplete clades”, and 24.7% are “Errors”. “Error” clades are larger (composed of more SNPs) than “Full clades” and “Incomplete clades”, but not “Multiple derived lineages” (*Mann-Whitney U)*. We further found that the average of the Jaccard Scores of SNPs that compose “Error” clades (∼0.11) tends to be lower than the other classifications, with approximately half (58/122) of the “Error” clades having a Jaccard Score of zero (Supplemental Figure 8c). This implies that a subset of lineage-informative SNPs in “Error” clades co-occur in insertions at a low frequency, but at a rate insufficient to drive a positive Silhouette Score. Therefore these clades may represent mergings of related lineages that share subsets of SNPs but are more diverged than “Multiple derived lineages”, and/or are closely related to other clades in the dataset. However, the clades with an average Jaccard Score of zero are likely true false-positive associations in clustering and represent ∼12% of all inferred clades (59 clades, 58 of which are “Error” clades).

### Calculating Sequence Diversity and Sequence Length of Clades using PacBio Genomes

We calculated the average sequence diversity of inferred clades using PacBio data by finding all insertions of a TE family that belonged to each clade from the PacBio alignments (only considering an insertion to belong to a clade if its sequence contained all detectable lineage-informative SNPs). From those insertions we calculated sequence diversity as Nei and Li (Nei and Li 1979), but when performing pairwise comparisons of nucleotide differences we only considered positions where the two sequences did not have gaps in either sequence.

We additionally calculated the average lengths of insertions belonging to each clade as a proportion of the full length consensus sequence. We called insertions belonging to a particular clade as above, counted the number of non-gap positions in each alignment, divided them by the total length of the TE consensus sequence and then averaged these proportions across all insertions within a clade.

### Distinguishing Heterochromatic and Euchromatic Insertions in PacBio Genomes

In order to determine whether a TE insertion in a PacBio genome is euchromatic or heterochromatic we first masked repeats in the PacBio genome assemblies and the release 6 *D. melanogaster* reference genome using RepeatMasker (Smit 2013). We next aligned the PacBio assemblies to the reference genome using *mummer 3.1* requiring a minimum 100 bps of alignment for each aligned portion (*nucmer -l 100 -p ${pacbio genome} ${reference genome}*) (Kurtz et al. 2004). We then took the coordinates of the pericentromeric heterochromatin boundary in the reference genome (Riddle et al. 2011) and found the corresponding position in the PacBio genome using the aligned segments of the genome. Exact matches to the reference genome heterochromatin boundary positions were difficult to obtain due to differences in assembly quality of the PacBio genomes, but we were typically able to find positions within 100 bp - 2 kb of the reference genome heterochromatin boundary coordinates that aligned uniquely to the PacBio genomes. The only exceptions were the X chromosome in N25 and the 2L arm of ZH26, where the closest uniquely aligned reference genome segments were 100-200 kb away from the coordinates of the heterochromatin boundary. Finally, we determined whether the position of TE insertions (taken from the TE alignments to the PacBio assemblies) were within heterochromatin or euchromatin, and recorded which clade each insertion belonged to as described in the above sections. We show the results of these data for all TE families except for telomeric TEs (Supplemental Figure 4c, 4d, 4e, 4f).

### Simulating Artificial Clades from Phylogenies

To generate the sequences of artificial clades, we first used a birth-death process to generate a topology of the evolutionary history of a TE (R package *treeSim) (Stadler 2011; Love et al. 2014)*. We reason that transposition events in a TEs evolutionary history can be considered births, while a deactivating mutation or excision would be equivalent to the extinction of a lineage. We used a birth rate of 1 × 10^-4^ and a death rate of 1 × 10^-5^(Le Rouzic et al. 2013). We simulated 20,560 generations of this process which generated a tree with 2,500 extant tips and 248 extinct tips -- a large but manageable number of sequences. We retained the extinct tips in the topology as they would represent TEs that are no longer active but still segregate in the population. We used this topology to generate sequences evolving neutrally by generating a random ancestral sequence of length 3,000 bp and dropping mutations via a Poisson process along the branch lengths with a mutation rate of 1 × 10^-7^ and no recombination, thus generating sequences for each tip (R package *simSeq*) (Schliep 2011). For a population of 85 individuals (the same number of individuals as in the GDL sample) we generated a copy-number distribution by drawing each individual’s copy number from a Poisson distribution with a mean copy number of 25. We then used this distribution to randomly sample from all extant and extinct lineages with replacement (Supplemental File 5, https://github.com/is-the-biologist/TE_CladeInference).

### Simulating Truncated Elements

In order to simulate the distribution within an individual of 5’ truncated elements, such as in LINE-like retrotransposons, we first sampled the total number of copies, *CN*, of an artificial element and its sequences as described above. We next generated the distribution of lengths of these elements by parameterizing a truncated geometric distribution with a mean of *L* and a maximum length of 3,000bp and drew *CN* times from this distribution. We assigned length values, *l*_*i*_, to each artificial TE sequence in an individual and removed positions from the 5’ end of each sequence such that we were left with *CN* TE sequences each *l*_*i*_long. We generated simulations where the average length of the elements (*L)* were 300, 600, 1,000, 1,500, 2,250 and 2,700 base pairs (10%, 20%, 33%, 50%, 75% and 90% of the full length element, respectively)

### Simulations of Short-Reads from Artificial Clades

We aimed to simulate data that would be obtained from short-read libraries generated genomes harboring TE insertions that were aligned to a consensus sequence using ConTExt. We used arrays of known sequences that “reside” in each simulated individual in our population to generate TE copy number, and an allele proportion matrix.

TE copy number is simply the number of copies of an artificial TE that an individual has within their “genome”. The allele proportion matrix contains the proportion of artificial TE sequences that contain an A, T, C, or G at a given position for each strain plus pseudocounts added to represent sequencing errors and mapping errors. We simulated error rates of 0%, 0.1%, 0.5%, 0.75%, 1%, 2.5% and 5%.

We then used these two reference files to simulate allele copy number pileups that replicate the inputs we used for our analysis of the GDL short-read data. For each strain we generated the coverage of our simulated library by drawing from a Poisson distribution with a target coverage as our lambda parameter:

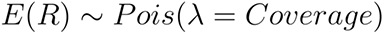

We call the values drawn from this distribution our Expected Reads, E(R). We then use E(R) to generate the observed number of reads, O(R), that map to a given position of our TE. We reason that the number of reads observed at a position would be the E(R) multiplied by the TE copy number, CN. Therefore, we draw O(R) from another Poisson distribution where lambda is E(R) times the CN:

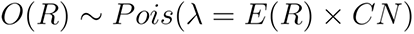

In the cases of truncated elements, where the copy number varies across the length of the element, we multiply E(R) by the copy number at each position to generate O(R). With the O(R) obtained for all positions of a TE for a given simulated library we now will use this to estimate the observed copy number, O(CN). We do this by adapting methods of copy number estimation, but instead of estimating E(R) with library specific parameters, we use our known E(R) from our simulated library (McGurk et al. 2020 Dec 22). In short we divide O(R) by E(R) to obtain our O(CN):

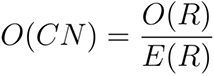

We now randomly sample O(R) number of reads from a multinomial distribution parameterized by the allele proportion matrix, thereby generating read counts that map to A, T, C or G. We use the proportion of reads that map to each nucleotide to generate a mapped allele proportion matrix that we multiply to O(CN) to obtain the number of copies observed for each allele. This was output as our final simulated allele copy-number matrix where we have recorded the copy number of each allele of a TE for 85 simulated libraries (Supplemental File 5, https://github.com/is-the-biologist/TE_CladeInference). These simulated data are identical in structure to the data structure that was used to infer clades from the GDL short-read data. We used our clade inference pipeline described in the above sections to infer clades using the same population specific parameters as the GDL (*π* > 0.1, Population Frequency > 10%).

### Simulations of TE copy-number data to benchmark clade inference

We benchmarked the performance of our clade inference method using the aforementioned simulations of sequence evolution and truncations to create artificial TE sequences segregating in a simulated population. We used the artificial sequences as a validation set for the clade inferences by calculating the frequency of inferred clades in the validation set and Silhouette Scores for each clade. When a set of parameters produced no interpretable data, *i.e.* no clusters were called, or a singular cluster encompassing all SNPs was called, we assigned a Silhouette Score of -1. We explored the parameter space of sequence errors, truncations of elements, and clustering cut-offs as well as their interactions to find which combinations of parameters led to lower Silhouette Scores when inferring clades.

First, we examined the interaction of sequencing error and clustering correlation cut-offs by generating simulated data where the sequencing error varied between 0% and 5% and clustering cut-off varied between 0 and 1, but all elements were full length (Supplemental Figure 2a). We find that even sequencing error rates as high as 5% (50x higher than Illumina sequencer error) have very little effect on clustering quality under most correlation cut-offs. This is likely because erroneous base calls are not incorporated downstream in the pipeline due to our SNP filtering steps, demonstrating the necessity of pre-processing the data for these types of analyses. Only when the correlation cut-offs were at extremes do we see a noticeable drop in the Silhouette Score.

We next examined the interaction between the average length of elements and correlation cut-offs when sequencing error was 0.1% (Illumina sequencing error rate). We randomly generated truncated elements with average lengths between 300bps and 2750bps (10 - 90% of the full length element) and found that truncations significantly decrease the ability to call clusters (Supplemental Figure 2b). Truncations and correlation cut-offs seem to interact negatively, wherein no clusters are called (Silhouette Score = -1) when the stringency is high and truncations are abundant. In the most extreme case (300bp elements) no positive correlations between alleles are found. This is an important consideration for highly fragmented TEs such as I-element, where many clades were validated but had negative Silhouette Scores, meaning that the fragmented nature of the TE broke down the positive correlations between SNPs and split clades apart.

Finally, we examined whether increased sequencing error and truncations interacted with each other at a set correlation cut-off, performing simulations using all pairwise combinations of truncation lengths and error rates described above (Supplemental Figure 2c). As might be expected, truncations and errors have compounding effects, where high error rates and high rates of deletion produce clusters of poorer quality than each parameter independently. Simulations with highly truncated elements and high error rates also tended to produce large numbers of clusters, occasionally in the thousands, which drove down clustering quality (Supplemental Figure 2d). This is an important interaction to consider as highly fragmented TEs may also have an increased rate of errors due to mappability of reads.

These simulations allowed us to generate a realistic dataset to benchmark our method and explore the parameter space of sequencing errors, deletions and correlation cut-offs that would negatively affect our inferences. A biological dataset will have some combination of deletions and sequencing errors depending on the TE, sequencing platform and alignment algorithm used, and choosing the optimal correlation cut-off should take the above information into account. However, given that many combinations of parameters produced positive Silhouette Scores and a relatively uniform number of clusters, except at extreme parameters, we are confident that our method can produce interpretable biological results even at non-optimal correlation cut-offs.

### Processing and aligning small RNA data

Public piRNA libraries were all created from female *D. melanogaster* ovaries, and are available through the SRA (see SRA accessions). We obtained libraries from 10 GDL strains (two from each population) (Luo et al. 2020). piRNA reads were trimmed using *Trimmomatic* and aligned to an index of curated RepBase repeat consensus sequences using *Bowtie2* with the parameters: *-N 1 -L 10 -i S,1,0.5 -p 8 --score-min L,0,-1.2 -D 100 -R 5* (Langmead and Salzberg 2012; Bolger et al. 2014; Bao et al. 2015; Langmead et al. 2019; McGurk et al. 2020 Dec 22). After alignment, reads were filtered by base quality (Q > 30), by size (21-30 base pairs) and by mapping quality as described for the genomic data in the above sections. From the remaining reads we generate SNP read pileups with the python module *pysam* using the *pileup* function (https://github.com/pysam-developers/pysam), akin to *samtools mpileup* (Li et al. 2009). We separated reads by sense and antisense to get SNP pileups derived from the secondary piRNA pathway, and the primary piRNA pathway, respectively. The result is a matrix containing the number of sense and antisense reads that map to each position and each read’s nucleotide at that position. After generating the matrices, we used a Size Factor Normalization approach to normalize the total read depth of all repeats that reads were aligned to. We generated a table of read counts for each TE from the SNP pileups, and then followed the protocols described by DESeq2, but used a custom script to handle our unique data structure (Love et al. 2014). The normalized read depth SNP pileups were used as the primary data for all piRNA analyses in this study. To calculate piRNA read depth of each clade we averaged the sense and antisense piRNA read depth across all alleles of each clade across the strains. We then added pseudocounts of one to the clade piRNA read depth and to the clade copy number of the strains before computing piRNAs/copy. This was done to regularize data for log-transformation. We used these values to calculate the average sense and antisense piRNA read depth per clade copy across the 10 GDL strains.

### Data and code availability

Short-read, PacBio and piRNA data used in this study were previously published and are publicly available: GDL NGS libraries are available under SRA accession SRP050151 (Grenier et al. 2015). GDL PacBio genomes are available under SRA accession SRP142531 (Long et al. 2018), and DSPR PacBio genomes are available under BioProject accession PRJNA418342 (Chakraborty et al. 2019). GDL piRNA data is available under SRA accession SRP068882 (Luo et al. 2020). Supplemental materials, code and processed data used to infer clades are publicly available through Github (https://github.com/is-the-biologist/TE_CladeInference).

## Acknowledgements

We thank Clayton Hughes for help with the piRNA alignments and Satyam Srivastav for invaluable discussion on the epigenetic inheritance of piRNA clusters. This work was supported by the General Medical Sciences Institute of the National Institutes of Health under NIGMS- R01-119125 to D.A.B. and A.G.C.

---

**Supplemental Figure 1.**
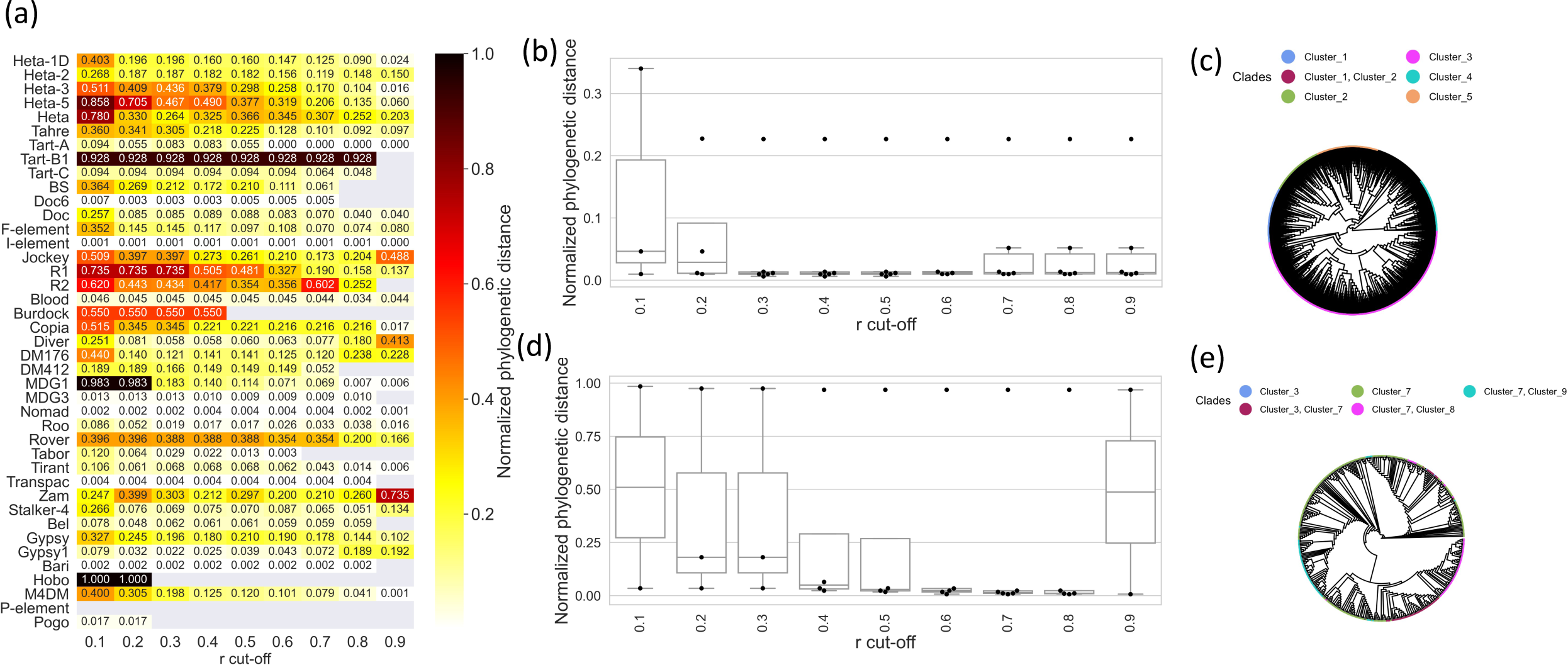
Relationship between clustering correlation cut-off and phylogenetic distance of clades. (a) Family-wise average of phylogenetic distance of tips belonging to each clade normalized by the sum of total branch lengths of the tree under different clustering correlation cut-offs of 41 TE families. Tips were determined to belong to a clade if their sequence contained two or more lineage-informative SNPs. Grey squares indicate no clusters were called or lineage-informative SNPs were not found in any sequence on the phylogeny. Phylogenies were constructed from TE insertions from PacBio genomes that were 75% full length or greater. (b) Phylogenetic distance of tips belonging to each clade normalized by the sum of total branch lengths of the tree under different correlation cut-offs for simulated TE data (Sequence error rate = 0.1%; all full length elements). Each point represents an inferred clade. (c) Phylogeny of simulated TE sequence data (Sequence error rate = 0.1%; all full length elements) with tips annotated by which clades they belong to under a correlation cut- off of *Pearson’s r* = 0.6. (d) Phylogenetic distance of tips belonging to each clade normalized by the sum of total branch lengths of the tree under different correlation cut-offs for the *Jockey* element. Each point represents an inferred clade. (e) Phylogeny of *Jockey* insertions from PacBio genomes with tips annotated by which clades they belong to under a correlation cut-off of *Pearson’s r* = ∼0.55.

**Supplemental Figure 2.**
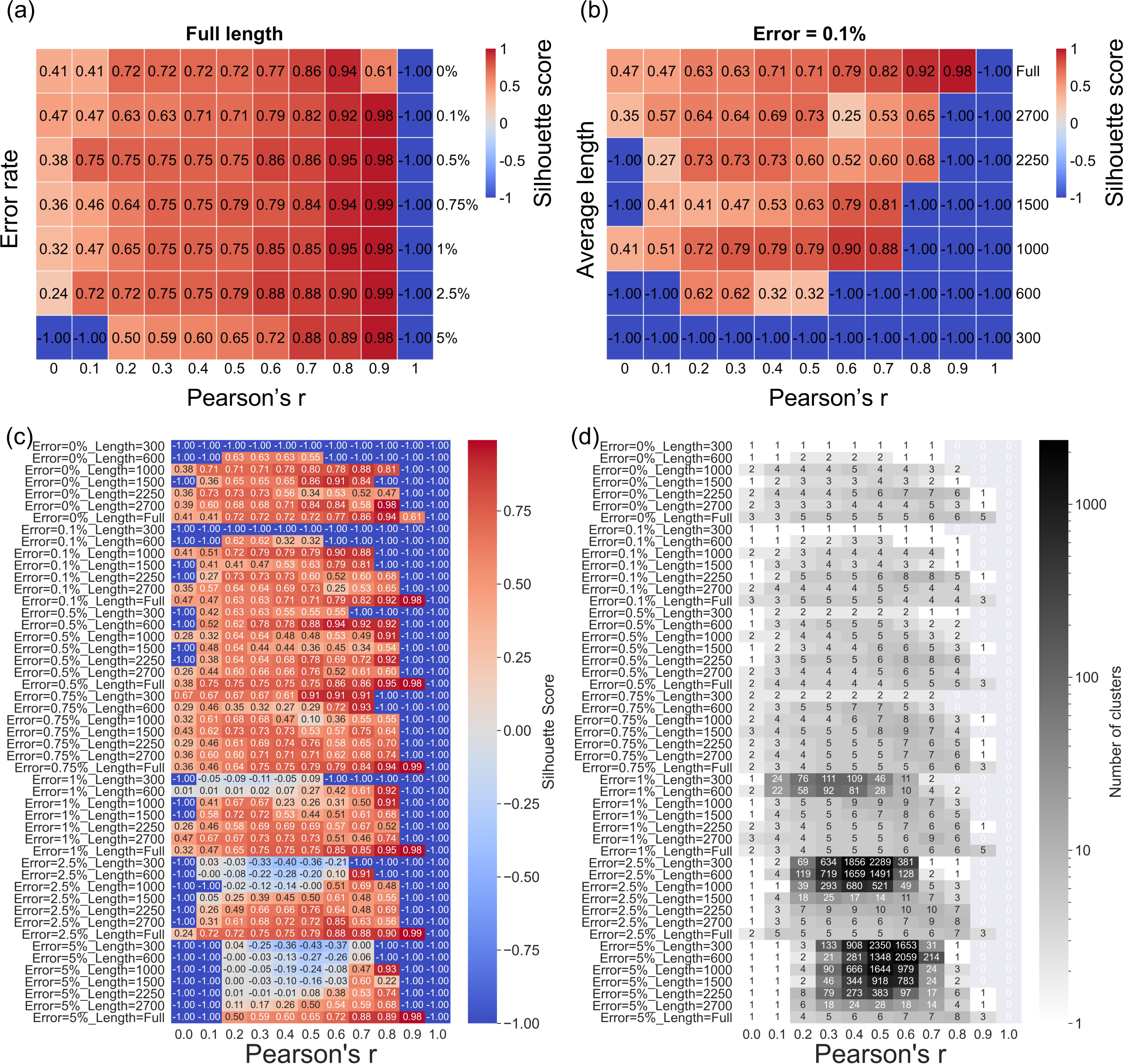
Benchmark of clade inference using simulated data. (a) Silhouette Score (a metric of clustering quality bound between -1 and 1, where low scores mean that SNPs are found more often in different elements than together, while high scores mean the opposite) results from inferring clades from simulated TE copy number data at variable sequence error rates and correlation cut-offs, where all sequences are full length. (b) Silhouette Score results from inferring clades from simulated TE copy number data at variable average length of elements (“Full” indicates only full-length elements) and correlation cut-offs, at a uniform sequence error rate of 0.1%. (c) Silhouette Score results from inferring clades from simulated TE copy number data using all pairwise combinations of sequencing error, average length of elements and correlation cut-offs (“Full” indicates only full-length elements). (d) Number of clusters inferred using simulated TE copy number data using all pairwise combinations of sequencing error, average length of elements and correlation cut-offs (“Full” indicates only full-length elements).

**Supplemental Figure 3.**
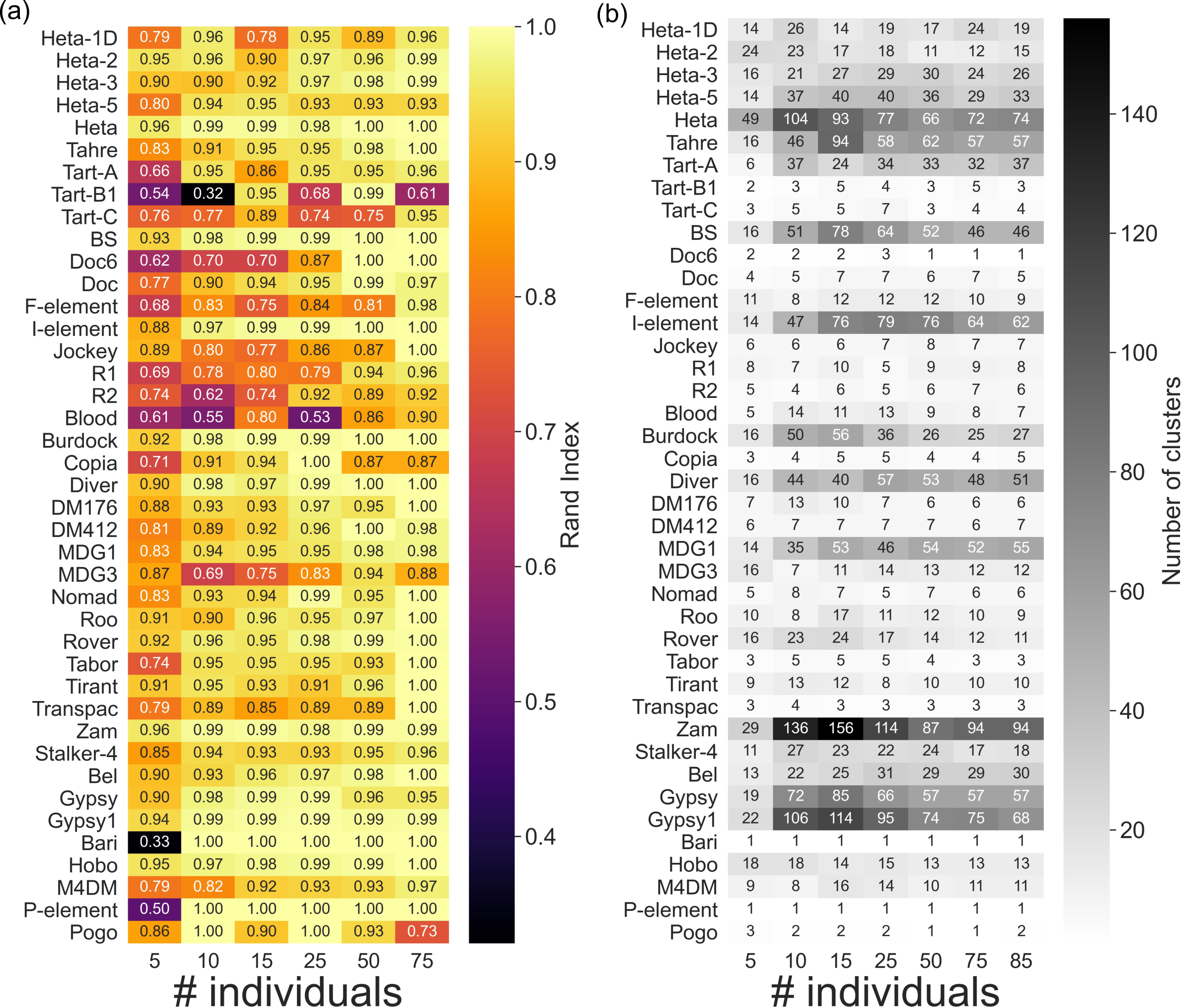
Effect of downsampling strains in clade inference. (a) Rand Index (proportion of accurate clusterings) of clade inference when downsampling the number of strains used to perform inference for all 41 TEs. (b) Number of clusters of clade inference when downsampling the number of strains used to perform inference for all 41 TEs.

**Supplemental Figure 4.**
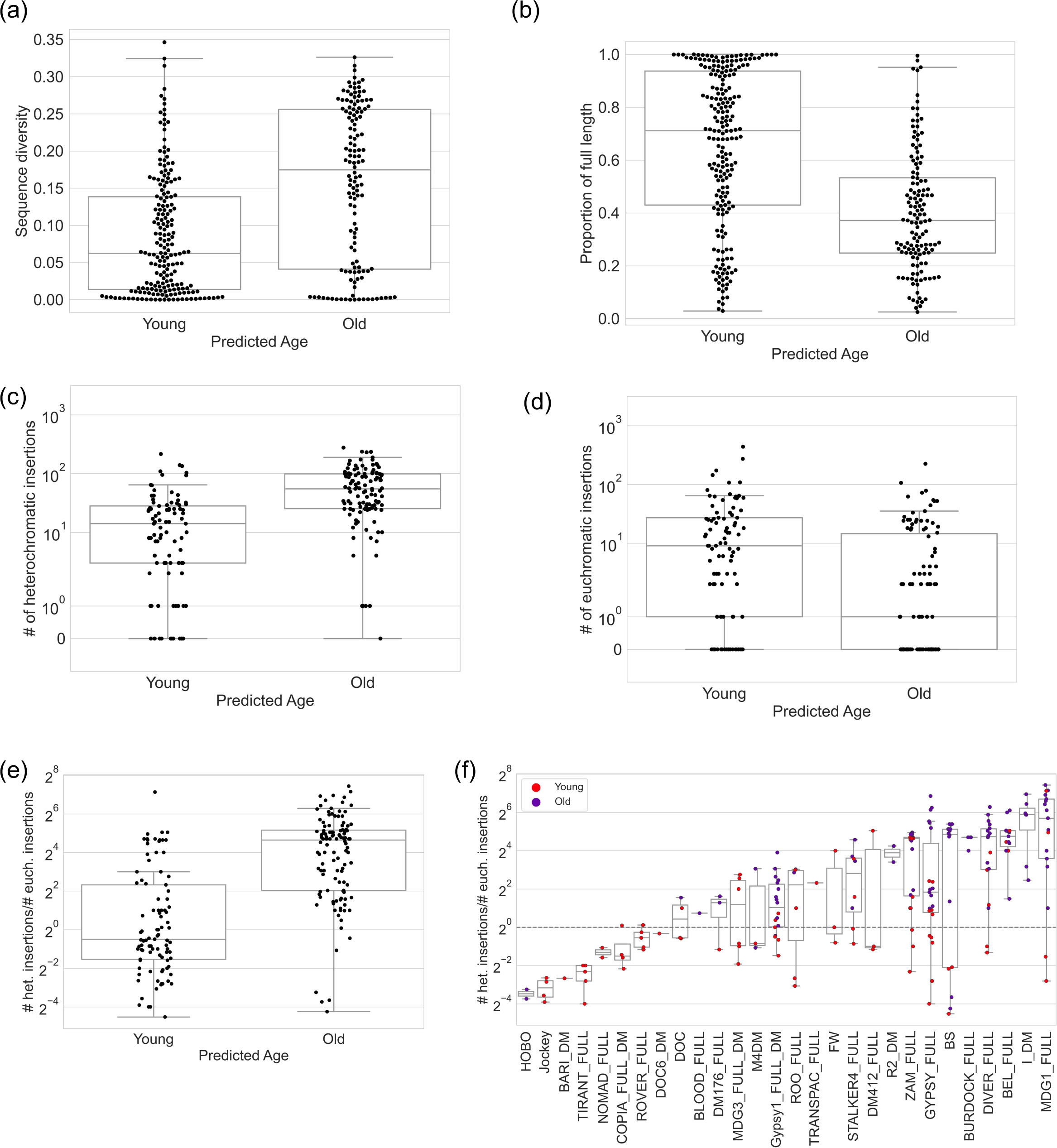
Validation of predicted age of inferred clades using PacBio data. (a) Sequence diversity of insertions in PacBio data belonging to young and old clades. (b) Average length of insertions in PacBio data belonging to young and old clades as a proportion of the full length consensus sequence. (c) Number of heterochromatic insertions in PacBio data belonging to young and old clades (excluding telomeric TE clades). (d) Number of euchromatic insertions in PacBio data belonging to young and old clades (excluding telomeric TE clades). (e) Ratio of heterochromatic to euchromatic insertions in PacBio data belonging to young and old clades (excluding telomeric TE clades). When computing the ratio of the number of heterochromatic and euchromatic insertions we regularized by adding 1 to the numerator and denominator. (f) Ratio of heterochromatic to euchromatic insertions in PacBio data belonging to each clade separated by TE family (excluding telomeric TE clades) and colored by predicted age (young; red, old; blue). When computing the ratio of the number of heterochromatic and euchromatic insertions we regularized by adding 1 to the numerator and denominator.

**Supplemental Figure 5.**
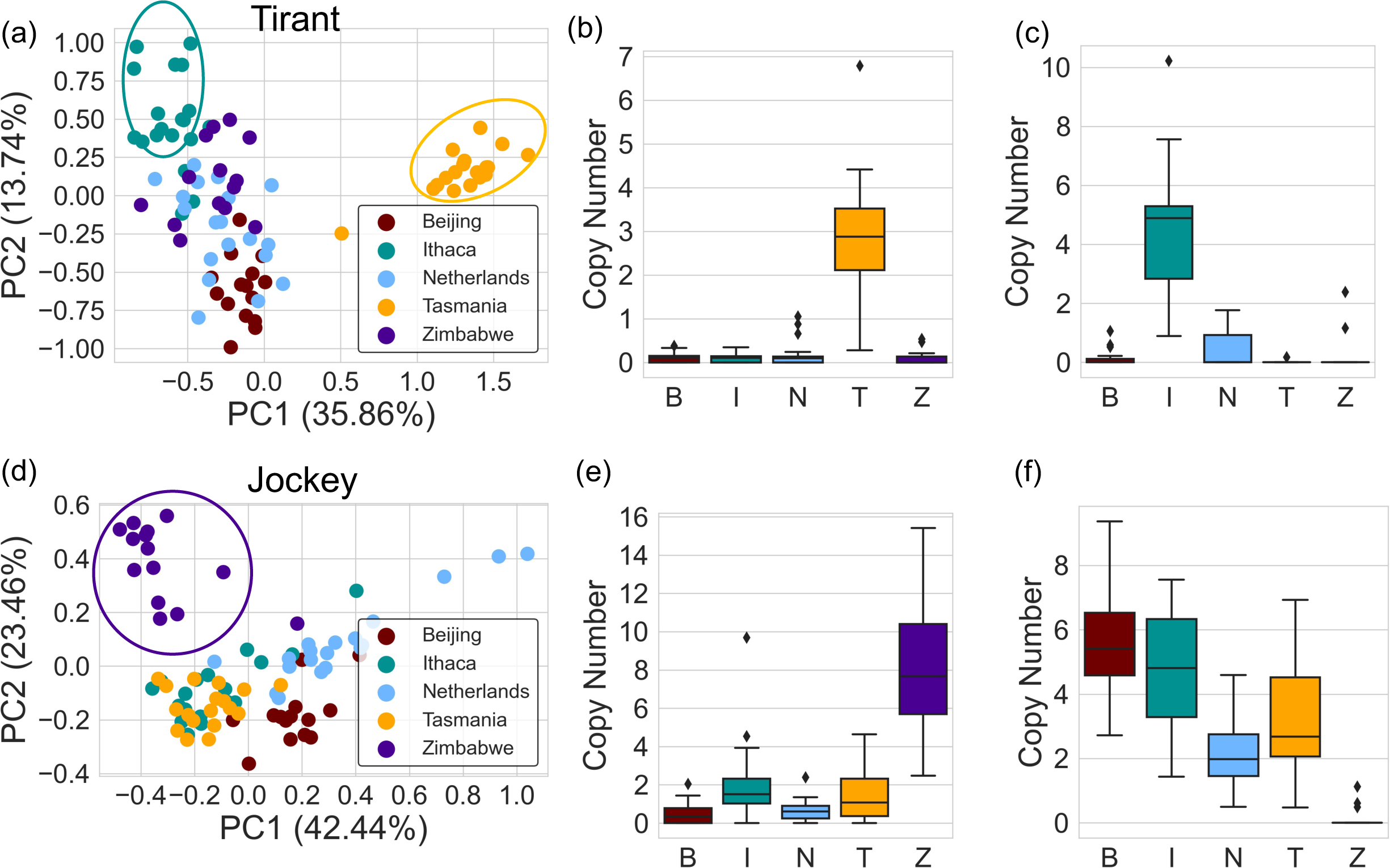
PCAs of minor allele frequencies from TEs that show strong population structure. (a) PCA on the minor allele frequency of *Tirant* elements reveals an Ithaca cluster (teal) and a Tasmania cluster (orange). (b) Boxplot of clade copy number of the *Tirant* clade enriched for Tasmania (B: Beijing, I: Ithaca, N: Netherlands, T: Tasmania, Z: Zimbabwe). (c) Boxplot of clade copy number of a *Tirant* clade enriched strongly for Ithaca, and moderately for Netherlands (B: Beijing, I: Ithaca, N: Netherlands, T: Tasmania, Z: Zimbabwe). (d) PCA on the minor allele frequency of *Jockey* elements reveals a cluster for Zimbabwe (purple). (e) Boxplot of clade copy number of a *Jockey* element enriched for Zimbabwe (B: Beijing, I: Ithaca, N: Netherlands, T: Tasmania, Z: Zimbabwe). (f) Boxplot of clade copy number of a *Jockey* element depleted in Zimbabwe (B: Beijing, I: Ithaca, N: Netherlands, T: Tasmania, Z: Zimbabwe).

**Supplemental Figure 6.**
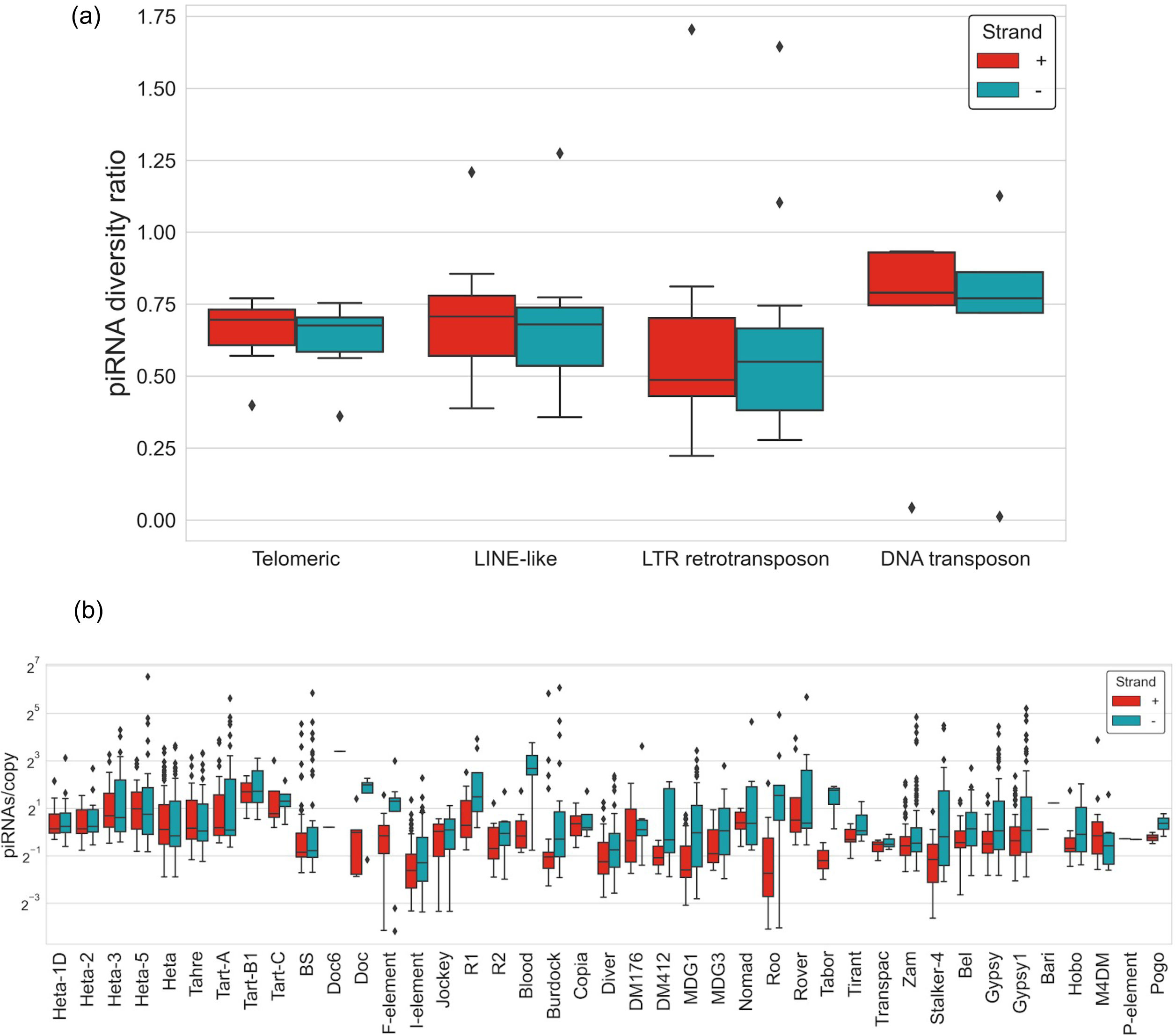
Additional piRNA diversity ratio and piRNA abundance data for recently active TEs. (a) Sense (red) and antisense (blue) piRNA diversity ratio (***π***_piRNA_/***π***_TE_) for all TE families from 10 GDL strains separated by class. (b) Average clade piRNAs/copy for sense (+, red), and antisense (-, blue) separated by TE family.

**Supplemental Figure 7.**
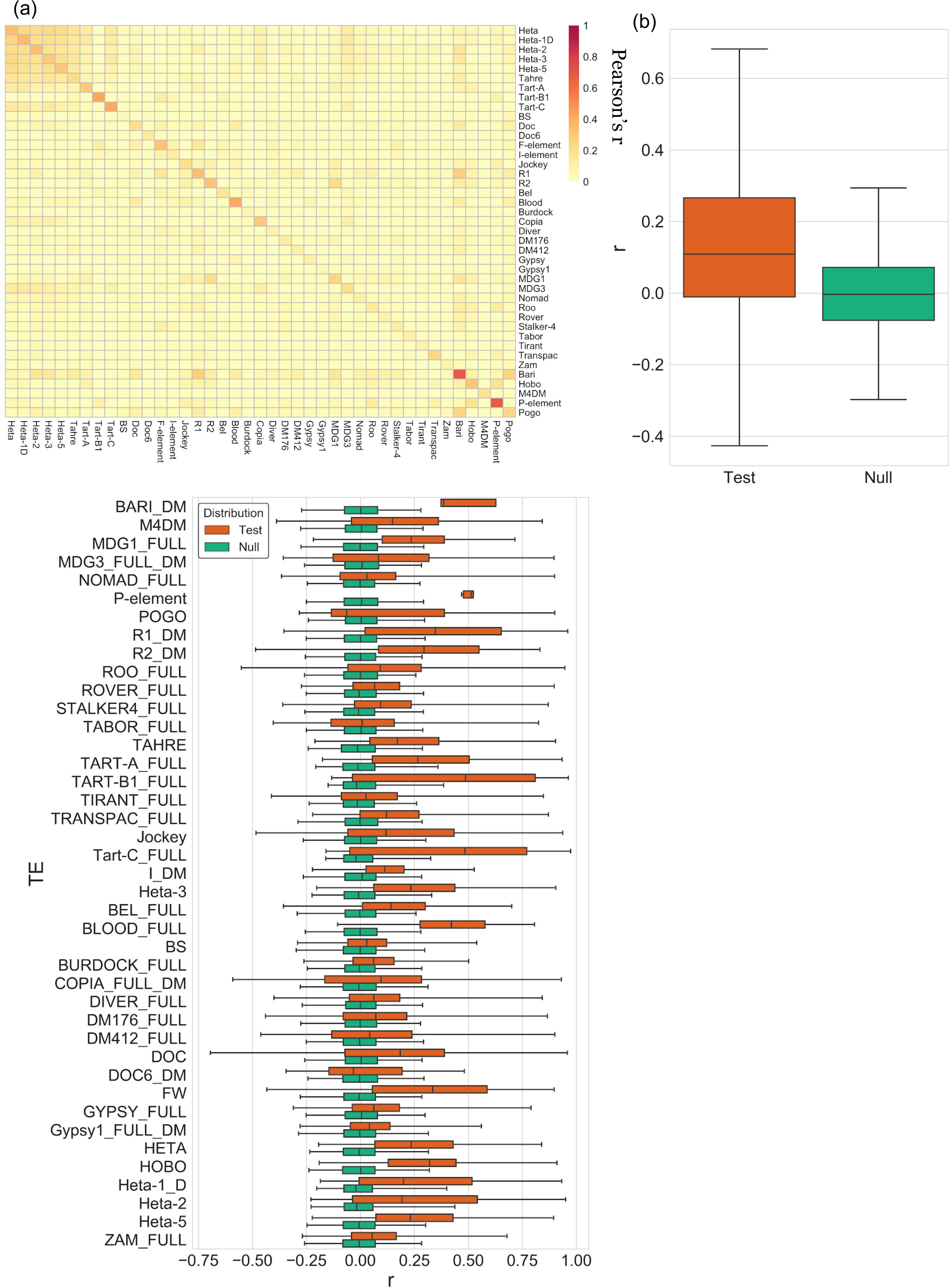
Correlations of SNPs within and between TE families and permutations used to choose optimal correlation cut offs for hierarchical clustering. (a) Average pairwise correlation of allele copy number between TE families and within TE families from GDL short-reads. Null distributions were constructed by permuting the allele copy number matrices for each TE family 1,000 times and recording all pairwise correlations between permuted alleles. The test distributions were computed by performing the pairwise comparisons of the allele copy number matrices for each TE family. (b) Aggregations of all test distributions (orange) and null distributions (teal) for all TE families. (c) Boxplots of individual test distributions (orange) and null distributions (teal) for each TE family.

**Supplemental Figure 8.**
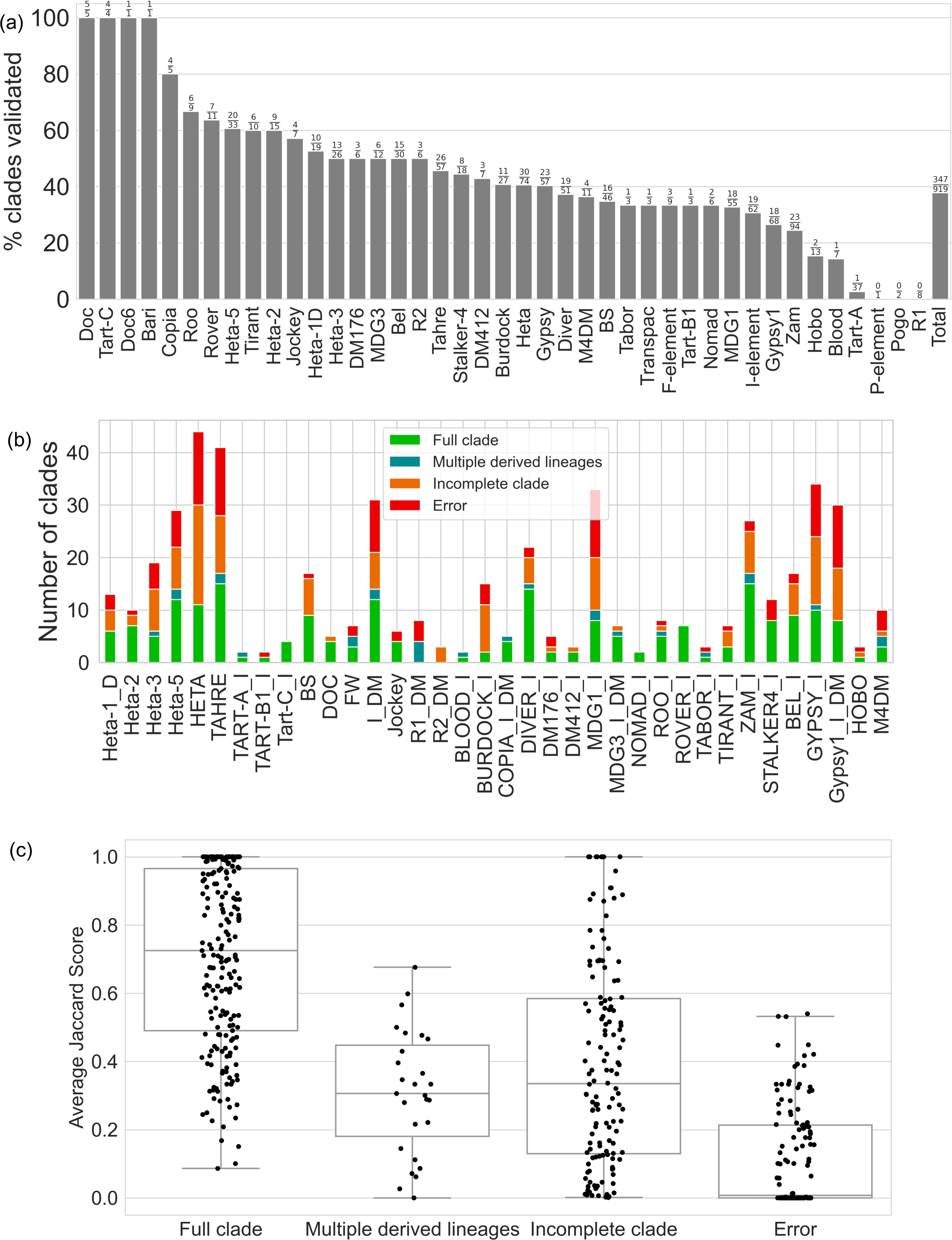
Additional validation of GDL clades using PacBio data. (a) The percent of all clades inferred from GDL data that were then detected in a set of PacBio genomes (including those with no SNPs detected in the PacBio data). The results are separated by TE family, with total clades shown on the far right. (b) Each clade is classified by a Silhouette Score and insertion frequency in the PacBio genomes: Full clade (Silhouette Score > 0, Frequency > 0; good clustering quality and detected in PacBio; green), Incomplete clade (Silhouette Score =< 0, Frequency > 0; poor clustering quality and detected in PacBio; orange), Multiple derived lineages (Silhouette Score > 0, Frequency = 0; good clustering quality and not detected in PacBio; blue), and Errors (Silhouette Score =< 0, Frequency = 0; poor clustering quality and not detected in PacBio; red). Results are separated by TE family. (c) Average Jaccard Score of each clade calculated as the average of the pairwise Jaccard Scores between all combinations of lineage-informative SNPs using PacBio genomes. Results separated by Silhouette Score classification and insertion frequency: Full clade (Silhouette Score > 0, Frequency > 0; green), Incomplete clade (Silhouette Score =< 0, Frequency > 0; orange), Multiple derived lineages (Silhouette Score > 0, Frequency = 0; blue), and Errors (Silhouette Score =< 0, Frequency = 0; red).

